# The protease ADAMTS5 controls ovarian cancer cell invasion, downstream of Rab25

**DOI:** 10.1101/2024.07.08.602517

**Authors:** Shengnan Yuan, Rachele Bacchetti, Jamie Adams, Elena Rainero

## Abstract

Ovarian cancer is the 3^rd^ most common gynaecological malignancy worldwide, with a 5-year survival rate of less than 30% in the presence of metastasis. Metastatic progression is characterised by extensive remodelling of the extracellular matrix, primarily mediated by secreted matrix metalloproteinases, including members of the ‘a disintegrin and metalloprotease with thrombospondin motif’ (ADAMTS) family. In particular, ADAMTS5 has been reported to be upregulated in ovarian malignant tumours compared to borderline and benign lesions, suggesting it might play a role in metastatic progression. Furthermore, it has been suggested that Rab25, a small GTPase of the Ras family, might upregulate ADAMTS5 expression in ovarian cancer cells. Here we demonstrated that Rab25 promotes ADAMTS5 expression, through the activation of the NF-κB signalling pathway. Furthermore, ADAMTS5 was necessary and sufficient to stimulate ovarian cancer cell migration through complex fibroblast-secreted matrices, while ADAMTS5 inhibition prevented ovarian cancer spheroid invasion in 3D systems. Finally, in ovarian cancer patients high ADAMTS5 expression correlated with poor prognosis. Altogether, these data identify ADAMTS5 as a novel regulator of ovarian cancer cell migration and invasion, suggesting it might represent a novel therapeutic target to prevent ovarian metastasis.

## Introduction

Ovarian cancer (OC) is the 3^rd^ lethal gynaecological cancer worldwide [1]. Patients’ death is primarily due to late-stage diagnosis, metastasis and chemotherapy resistance [2]. While the 5-year survival rate of OC patients diagnosed at the early stages is over 70%, this drops to less than 30% in the presence of peritoneal or distant metastasis [3]. The peritoneum represents the primary metastatic site and metastatic colonisation is characterised by extensive remodelling of the extracellular matrix (ECM), a dynamic network of secreted proteins.

The tumour microenvironment (TME) and the ECM play a key role in promoting cancer development and metastasis [4]. The ECM is continuously remodelled by ECM-modifying enzymes, including matrix metalloproteinases, secreted by both cancer cells and cancer-associated fibroblasts (CAFs) [5]. Indeed, altered ECM remodelling, caused by deregulated expression of ECM-modifying enzymes, was found to promote the proliferation, migration and invasion of tumour cells from different cancer types, including OC [6].

A disintegrin and metalloprotease with thrombospondin motif (ADAMTS) is a family of 19 secreted zinc-dependent metalloproteases [7], the expression of which has been reported to be dysregulated in multiple cancer types [8]. In OC, ADAMTS5 expression was found significantly increased in malignant tumour samples compared to borderline and benign tumours, suggesting that ADAMTS5 could promote OC metastasis [9]. However, the role of ADAMTS5 in controlling OC cell migration and invasion is still unclear. Interestingly, ADAMTS5 has been suggested to be upregulated in OC cells over-expressing Rab25 [10], a small GTPase of the RAS superfamily [11], whose expression was found significantly upregulated in advanced OC stages [12]. At the molecular level, Rab25 promoted OC cell migration and invasion in 3D matrices by enhancing the recycling of the ECM receptor integrin α5β1 at the pseudopod tips of OC cells [13].

Here we demonstrated that Rab25 promoted the expression of ADAMTS5, in a nuclear factor κB (NF-κB)-dependent manner. Importantly, secreted ADAMTS5 was necessary and sufficient to drive OC cell migration through fibroblast-generated 3D matrices. Furthermore, Rab25 and ADAMTS5 downregulation opposed CAF-driven ovarian cancer cell spheroid invasion in 3D systems, without affecting cell proliferation. Finally, elevated ADAMTS5 expression correlated with poor prognosis in ovarian cancer patients. Altogether, these indicate that ADAMTS5 could represent a novel druggable therapeutic target to prevent OC migration and invasion.

## Results

### Rab25 induced ADAMTS5 expression in ovarian cancer cells

ADAMTS5 was identified in a microarray screen to detect ECM- and Rab25-dependent changes in gene expression in OC cells [10]. To validate the role of Rab25 in inducing ADAMTS5 expression, A2780 cells stably expressing Rab25 (A2780-Rab25) or empty vector pcDNA3 controls (A2780-DNA3) were seeded on plastic or cell-derived matrix generated by telomerase immortalised fibroblasts (TIF-CDM), a 3D matrix rich in collagen and fibronectin which recapitulates multiple features of ECM in *vivo* [14] and the expression level of ADAMTS5 was measured by RT-qPCR (**Figure 1A**). Remarkably, we detected a 2.5-fold increase in ADAMTS5 expression in A2780-Rab25 cells compared to A2780-DNA3 cells on CDM, while a smaller, but still statistically significant, increase (1.5-fold) was measured on plastic (**Figure 1B**). These data indicate that the Rab25 increased ADAMTS5 mRNA levels and the presence of ECM further enhanced Rab25-dependent ADAMTS5 expression in OC cells.

**Figure 1.**
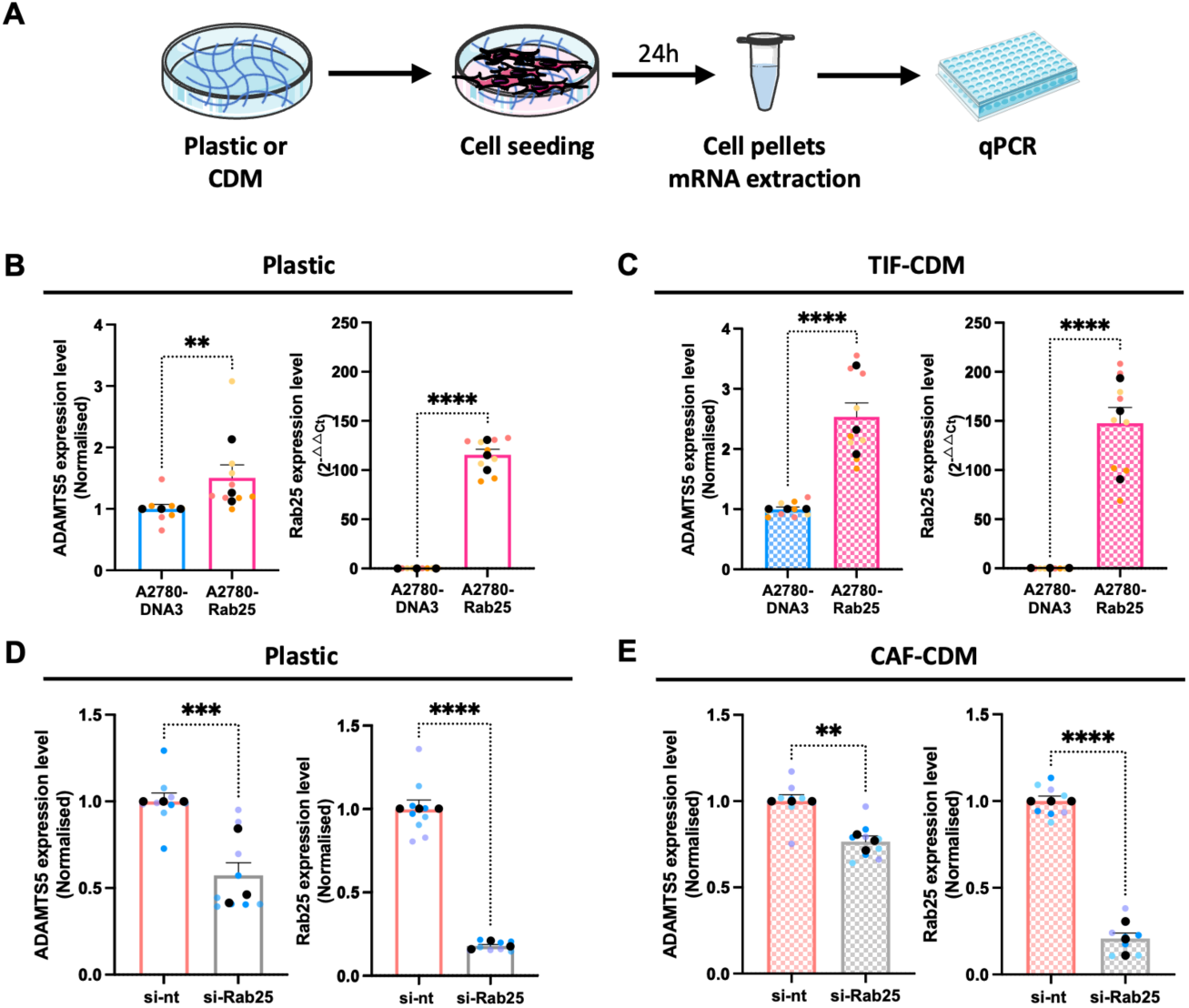
Rab25 induced ADAMTS5 expression in OC cells. **(A)** Schematic of the experimental plan. **(B,C)** A2780-DNA3 and A2780-Rab25 cells were seeded on plastic (B) or TIF-CDM (C) and the mRNA was extracted and quantified by qPCR. The data were normalised to A2780-DNA3 cells (ADATMS5) or plotted in 2 ^AACt^ (Rab25). Data are mean ± SEM from N=3 independent experiments. The black dots represent the mean of individual experiments. **p=0.005, ****p<0.0001, Mann-Whitney test. **(D,E)** OVCAR3 cells were transfected with a non-targeting siRNA control (si-nt) or a Rab25-targeting siRNA (si-Rab25) and seeded on plastic **(D)** or CAF-CDM **(E).** ADAMTS5 and Rab25 mRNA levels were quantified by qPCR and normalised to si-nt. Data are presented as mean ± SEM from N=3 independent experiments. The black dots represent the mean of individual experiments. **p=0.0012, ***p=0.0003, ****p<0.0001, Mann-Whitney test. Image created with items adapted from Servier Medical Art, licensed under CC BY 4.0.

Since ADAMTS5 is a secreted protease, to confirm the changes in ADAMTS5 expression at the protein level, conditioned media was harvested from A2780-DNA3 and A2780-Rab25 cells and analysed by Western Blotting (**Figure S1B**). Similar to the mRNA expression data, ADAMTS5 protein levels were significantly increased in A2780-Rab25 cells seeded on CDM, while there was no significant difference in ADAMTS5 levels on plastic (**Figure S1B**).

To study the role of Rab25 in a more physiological context, we measured endogenous Rab25 expression in a panel of OC cell lines and found that OVCAR3, but not OVCAR4 and SKOV3, overexpressed Rab25, to a similar level to the exogenous overexpression in A2780 cells (**Figure S1A**). OVCAR3 cells were transfected with a non-targeting siRNA control or an siRNA targeting Rab25 and ADAMTS5 expression was assessed by qPCR in cells seeded on plastic or CDM generated by omental CAFs [15]. Consistent with our previous data, the downregulation of Rab25 significantly reduced ADAMTS5 mRNA levels in OVCAR3 cells, both on plastic and on CDM, although here there was a more prominent effect on plastic (**Figure 1D,E**). These data demonstrate that Rab25 induced ADAMTS5 expression in OC cells, while the ECM had a more prominent role in A2780 compared to OVCAR3 cells.

### The transcription factor NF-κB was required for Rab25-induced ADAMTS5 expression in OC cells

Having established that Rab25 promoted ADAMTS5 expression, we set out to characterise the transcription factor(s) responsible for this. Rab25 has been reported to stabilise hypoxia inducible factor 1α (HIF1α) protein levels in normoxia in OC cells [16]. Since HIF1α was reported to weakly bind to ADAMTS5 promoter [17], we investigated the role of HIF1 in ADAMTS5 expression by treating the cells with a well-characterised HIF inhibitor, echinomycin [18], or incubating the cells under hypoxia for 24h. Surprisingly, echinomycin treatment significantly increased ADAMTS5 expression in both OVCAR3, endogenously over-expressing Rab25, and in SKOV3 cells, which lack detectable Rab25 expression (**Figure S2A-C**). Consistently, hypoxia incubation significantly reduced ADAMTS5 expression in all the cell lines tested, regardless of the expression of Rab25 (**Figure S2D-G**). Together, these data indicate that HIF1 is a negative regulator of ADAMTS5 expression and is not involved in Rab25-dependent ADAMTS5 expression in OC cells.

ADAMTS5 has been extensively studied in osteoarthritis, where its catalytic activity has been shown to contribute to the disease progression [19]. In chondrocytes, a subunit of transcription factor NF-κB, RelA/p65, was found to interact with ADAMTS5 promoter and strongly induce ADAMTS5 expression [17]. Therefore, we tested whether NF-κB also promoted ADAMTS5 expression in OC cells. BAY 11-7082 is a small molecule inhibitor specifically targeting IKK (inhibitor of nuclear factor kappa B kinase), which prevents the phosphorylation and degradation of IκBα (inhibitor of nuclear factor kappa B). As a result of BAY 11-7082 treatment, the release, nuclear translocation and promoter binding of NF-κB are inhibited (**Figure 2A**) [20]. OVCAR3 cells were treated with DMSO, 2.5, 5 or 10μM of BAY 11-7082 for 24h and the expression of ADAMTS5 was quantified by qPCR. As a result, ADAMTS5 expression decreased in a dose-dependent manner (**Figure 2B**). Importantly, Rab25 expression was not affected by BAY 11-7082 treatment (**Figure 2C**), indicating that NF-κB regulated ADAMTS5 expression downstream of Rab25.

**Figure 2.**
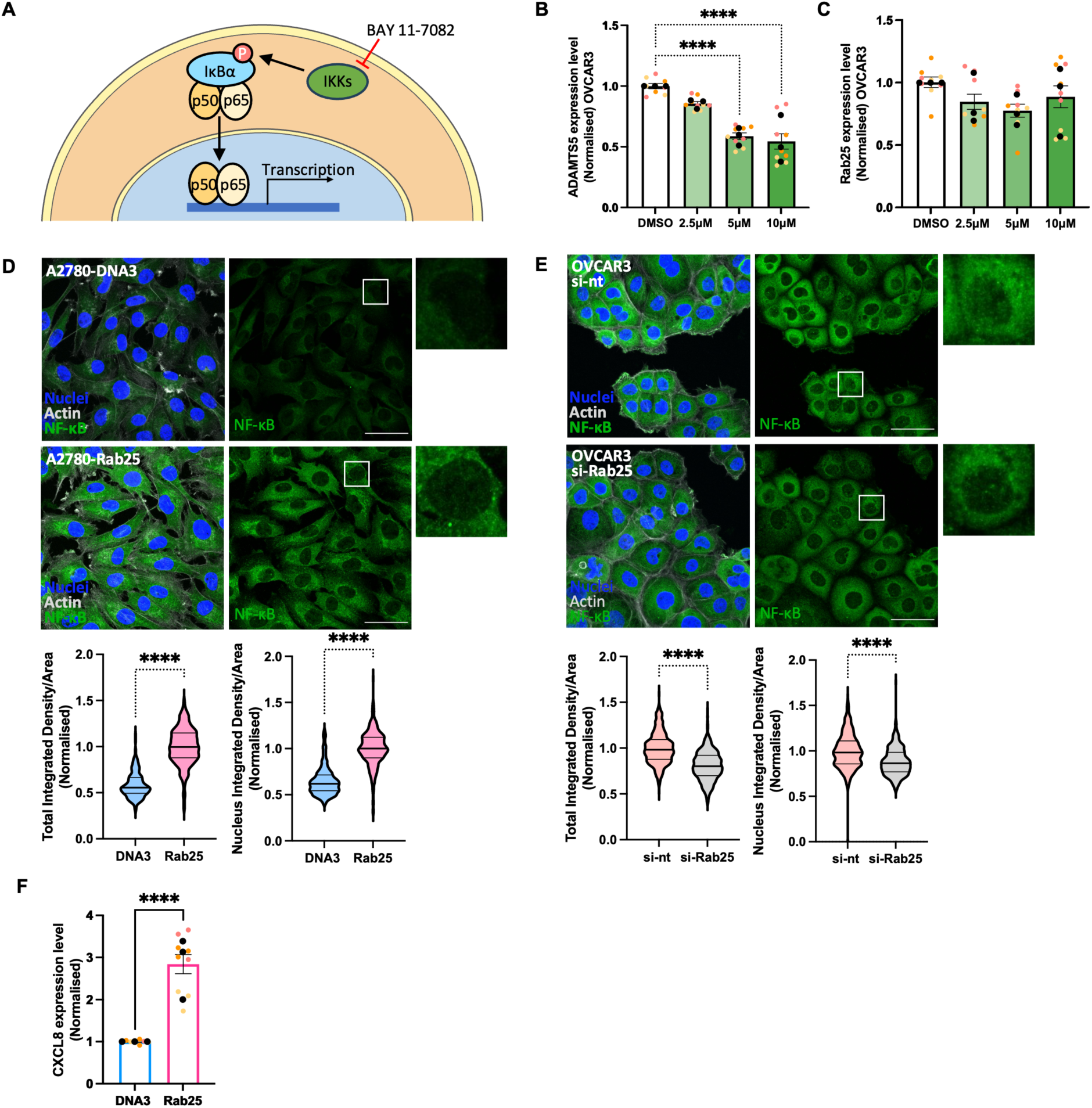
Rab25 induced ADAMTS5 expression in an NF-kB pendent-manner. **(A)** Schematic, NF-kB signalling pathway. **(B,C)** OVCAR3 cells were treated with DMSO, 2.5, 5 or 10μM of BAY 11-7082 for 24hr and the mRNA levels of ADAMTS5 (B) and Rab25 **(C)** were measured by qPCR. Data were normalised to the DMSO control and presented as mean ± SEM from N=3 independent experiments. The black dots represent the mean of individual experiments. ****p<0.0001, Kruskal-Wallis test. **(D,E)** A2780-DNA3, A2780-Rab25 cells (D), OVCAR3 cells transfected with non-targeting (si-nt) or Rab25 targeting (si-Rab25) si-RNA (E) were seeded on glass-bottom dishes, fixed, stained for NF-kB (green), nuclei (blue) and actin (grey), and imaged with a Nikon Al confocal microscope. Scale bar, 50μM. NF-kB integrated density for the whole cell and the nucleus was measured with image J and normalised to the cell area. Data were plotted as violin plots (median and quartiles) from N=3 independent experiments. ****p<0 0001, Mann-Whitney test. (F) CXCL8 mRNA levels in A2780-DNA3 and Rab25 cells were quantified by qPCR and normalised to A2780-DNA3 cells. Data are presented as mean ± SEM from N=3 independent experiments. The black dots represent the mean of individual experiments. ****p<0.0001, Mann-Whitney test.

To elucidate whether Rab25 controlled NF-κB signalling, we assessed the intracellular localisation of the NF-κB subunit RelA/p65 in A2780-DNA3 and A2780-Rab25 cells. RelA/p65 staining was predominately cytoplasmic, with weak nuclear staining. RelA/p65 signal intensity was significantly increased within the whole cells and in the nuclei of Rab25-overexpressing A2780 cells compared to DNA3 cells (**Figure 2D**). Similarly, Rab25 knockdown in OVCAR3 cells resulted in a small, but statistically significant, reduction in RelA/p65 overall and nuclear intensity, in comparison to the non-targeting control (**Figure 2E**). To confirm the role of Rab25 in promoting NF-κB activation, we measured the expression of CXCL8, a canonical NF-κB target gene, encoding for IL-8 [21]. Consistently, Rab25 over-expression in A2780 cells significantly increased CKCL8 mRNA levels (**Figure 2F**). Taken together, these results suggest that Rab25 induced ADAMTS5 expression through the regulation of NF-κB activation.

Several signalling pathways have been shown to control NF-κB. For instance, EGF-mediated EGFR stimulation was found to promote NF-κB activation via PI3K/AKT and MAPK/ERK pathways in OC cells [22]. Interestingly, Rab25 downregulation was found to impair EGFR signalling, with a consequent reduction in ERK and AKT signalling in radioresistant lung adenocarcinoma and nasopharyngeal carcinoma cells [23]. We therefore investigated whether Rab25 controlled these signalling pathways in OC cells. However, Rab25 overexpression in A2780 cells [24] or Rab25 knockdown in OVCAR3 cells did not affect AKT phosphorylation (**Figure S2A**). Similarly, Rab25 overexpression did not affect ERK phosphorylation in A2780 cells (**Figure S2B**), suggesting that Rab25 controlled NF-κB through an AKT- and ERK-independent signalling pathway.

### ADAMTS5 was required for Rab25-dependent OC cell migration on CDM

Rab25 was previously shown to promote OC cell migration in 3D matrices by enhancing integrin α5β1 recycling at the pseudopod tips [13]. We therefore wanted to investigate whether ADAMTS5 played a role in Rab25-dependent cell migration, by seeding A2780-DNA3 and A2780-Rab25 cells on CDM in the presence of DMSO control or 5μM ADAMTS5 inhibitor. Consistent with previous data [13], Rab25 overexpression significantly increased pseudopod elongation and directionality of cell migration, without affecting migration velocity. Remarkably, inhibition of ADAMTS5 catalytic activity significantly suppressed pseudopod elongation and directional migration in Rab25 overexpressing cells, without changing the morphology and migration of A2780-DNA3 cells (**Figure S3A-C**). Importantly, Rab25 overexpression and ADAMTS5 inhibition did not affect cell migration on plastic (**Figure S3D-F**), indicating that Rab25 and ADAMTS5’s role in controlling OC cell migration was ECM-dependent. Similarly, ADAMTS5 knockdown in A2780-Rab25 cells significantly reduced pseudopod length and directional cell migration on CDM (**Figure S3 G-J**). Together, these data demonstrate that ADAMTS5 was required for Rab25-dependent cell migration on CDM.

To more directly characterise the role of secreted ADAMTS5 in OC cell migration, we incubated A2780-DNA3 cells with conditioned media generated by A2780-Rab25 and DNA3 cells (**Figure 3A**). Compared to A2780-DNA3 conditioned media, treatment of A2780-DNA3 cells with A2780-Rab25 conditioned media significantly increased pseudopod length and directionality of cell migration, while the migration velocity was not affected (**Figure 3B**). To confirm that ADAMTS5 catalytic activity was required for this, A2780-DNA3 cells were treated with A2780-Rab25 conditioned media in the presence of DMSO or 5μM ADAMTS5 inhibitor and their migration ability was measured (**Figure 3C**). As a result, ADAMTS5 inhibition reduced the pseudopod length and migration directionality of A2780-DNA3 cells treated with A2780-Rab25 conditioned media (**Figure 3D**). Interestingly, ADAMTS5 inhibition also reduced the migration velocity of A2780-DNA3 cells, which was not identified in the absence of conditioned media (**Figure S3C**).

**Figure 3.**
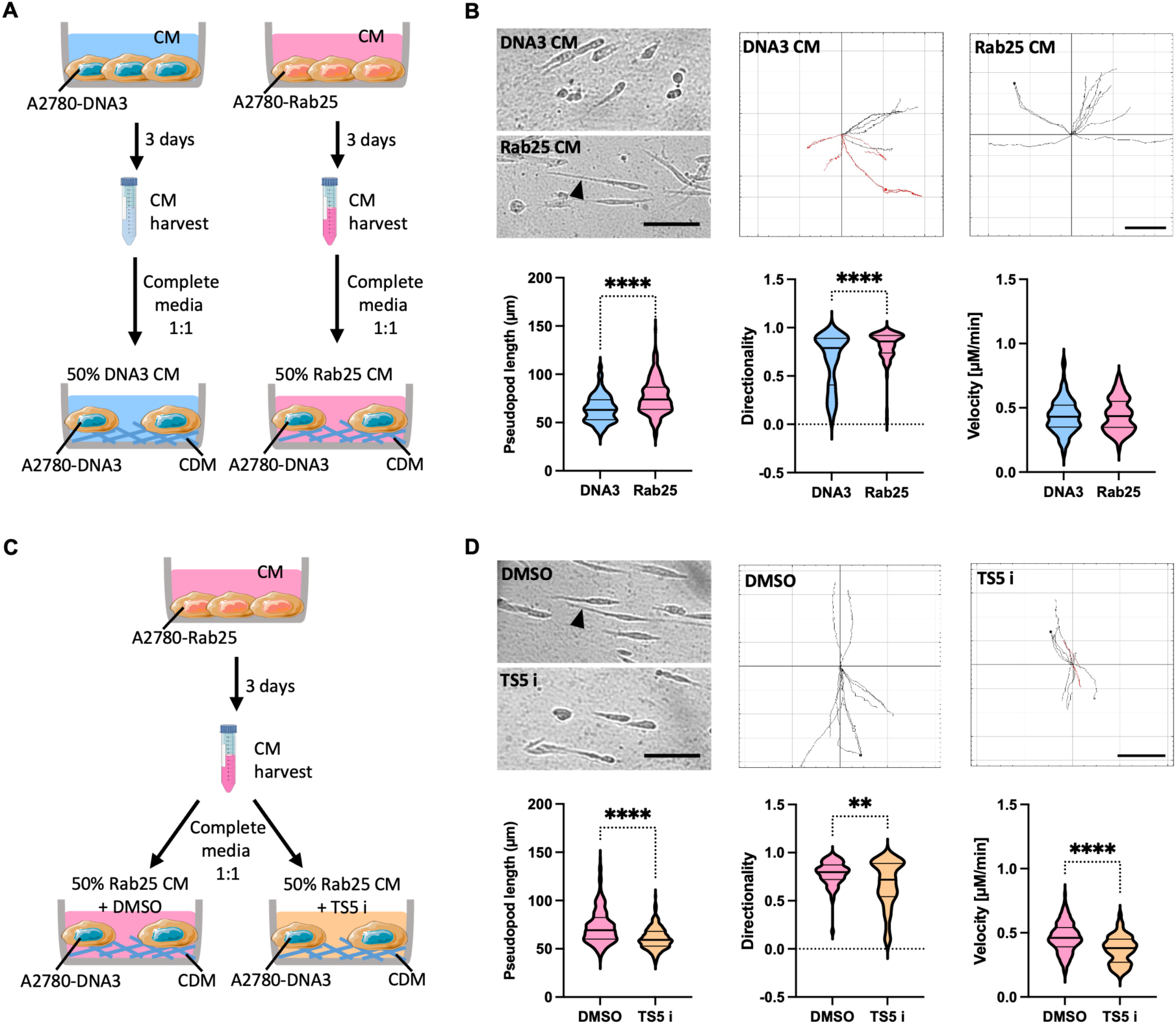
Conditioned media (CM) derived from A2780-Rab25 cells enhanced pseudopod elongation and directional migration of A2780-DNA3 cells on CDM, in an ADAMTS5-dependent manner. **(A,C)** Schematics of the experimental workflow. Conditioned media (CM) was collected from A2780-DNA3 and A2780-Rab25 cells seeded on plastic for 3 days. **(B,D)** A2780-DNA3 cells were seeded on CDM and treated with A2780-DNA3 or Rab25 CM (diluted 1:1 in complete media) (B), or with Rab25 CM in the presence of DMSO control or 5μM ADAMTS5 inhibitor (TS5 i, D). Cells were imaged live with a Nikon widefield live-cell system (Nikon Ti eclipse with Oko-lab environmental control chamber) for 16hr. Stills extracted from the movies are presented. The arrow-heads point to the elongated pseudopods. Scale bar, 100μM. Representative spider plots show the migration path of manually tracked cells (directionality >0.5 in black, <0.5 in red). Scale bar, 200μM. The pseudopod length (μm), directionality and velocity [μM/min] were measured with ImageJ. Data were plotted as violin plots (median and quartiles) from N=3 independent experiments. **p=0.0018, ****p<0.0001, Mann-Whitney test. Image created with items adapted from Servier Medical Art, licensed under CC BY 4.0.

Several factors, other than ADAMTS5, could be present in Rab25-overexpressing cell conditioned media and promote cell migration. Therefore, conditioned media was generated from control or ADAMTS5 knockdown A2780-Rab25 cells and added to A2780-DNA3 cells seeded on CDM (**Figure 4A**). Consistent with the ADAMTS5 inhibition results, A2780-DNA3 cells treated with conditioned media derived from ADAMTS5 knockdown A2780-Rab25 cells showed a significantly lower pseudopod length, migration directionality and velocity compared to conditioned media generated from non-targeting siRNA treated cells (**Figure 4B**). Altogether, these results demonstrate that secreted ADAMTS5 was sufficient to drive OC cells migration on CDM.

**Figure 4.**
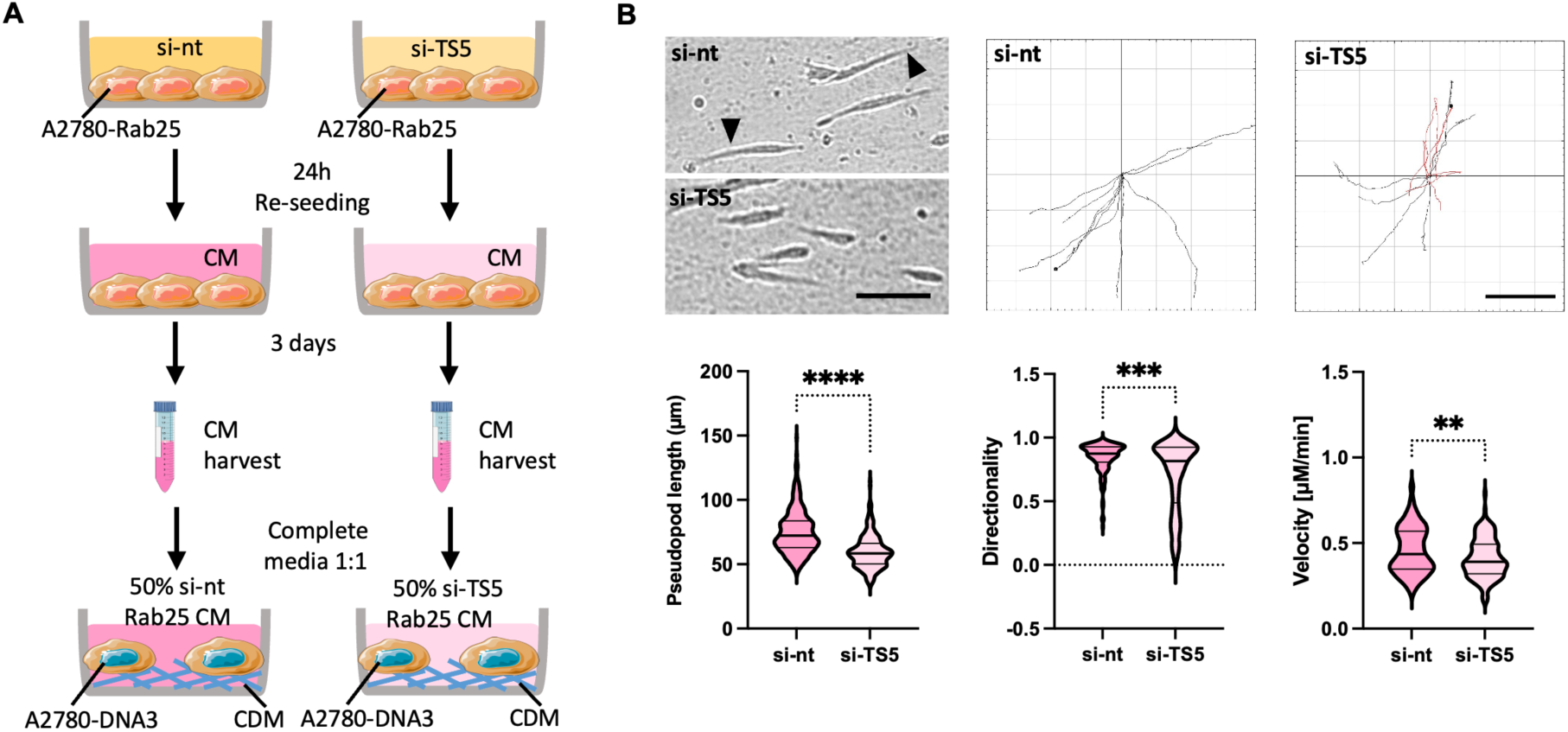
CM derived from ADAMTS5 KD A2780-Rab25 cells failed to promote the migration of A2780-DNA3 cells on CDM. **(A)** Schematic of the experimental workflow. Conditioned media (CM) was collected from A2780-Rab25 cells transfected with a non­targeting siRNA control (si-nt) or ADAMTS5 targeting si-RNA (si-TS5) and grown on plastic for 3 days. **(B)** A2780-DNA3 cells were seeded on CDM and treated with si-nt or si-TS5 A2780-Rab25 CM (diluted 1:1 in complete media). Cells were imaged live with a Nikon widefield live-cell system (Nikon Ti eclipse with Oko-lab environmental control chamber) for 16hr. Stills extracted from the movies are presented. The arrow-heads point to the elongated pseudopods. Scale bar, 100μM. Representative spider plots show the migration paths of manually tracked cells (directionality >0.5 in black, <0.5 in red). Scale bar, 200μM. The pseudopod length (μm), directionality and velocity [μM/min] of cell migration were measured with Imaged. Data were plotted as violin plots (median and quartiles) from N=3 independent experiments. **p=0.0057, ***p=0.0002, ****p<0.0001, Mann-Whitney test. Image created with items adapted from Servier Medical Art, licensed under CC BY 4.0.

As NF-κB was required for Rab25-induced ADAMTS5 expression, we tested the effect of NF-κB inhibition on the migration ability of A2780-Rab25 cells. The cells were pretreated with DMSO or BAY 11-7082 for 24h and cell migration on CDM was measured (**Figure 5A**). Compared to the DMSO control, treatment with BAY 11-7082 resulted in a dose-dependent reduction in pseudopod length and migration directionally, while the velocity of cell migration was not affected (**Figure 5B, C**). These results show that NF-κB signalling played a role in Rab25-dependent migration of OC cells, potentially by promoting the expression of ADAMTS5 (**Figure 2B**).

**Figure 5.**
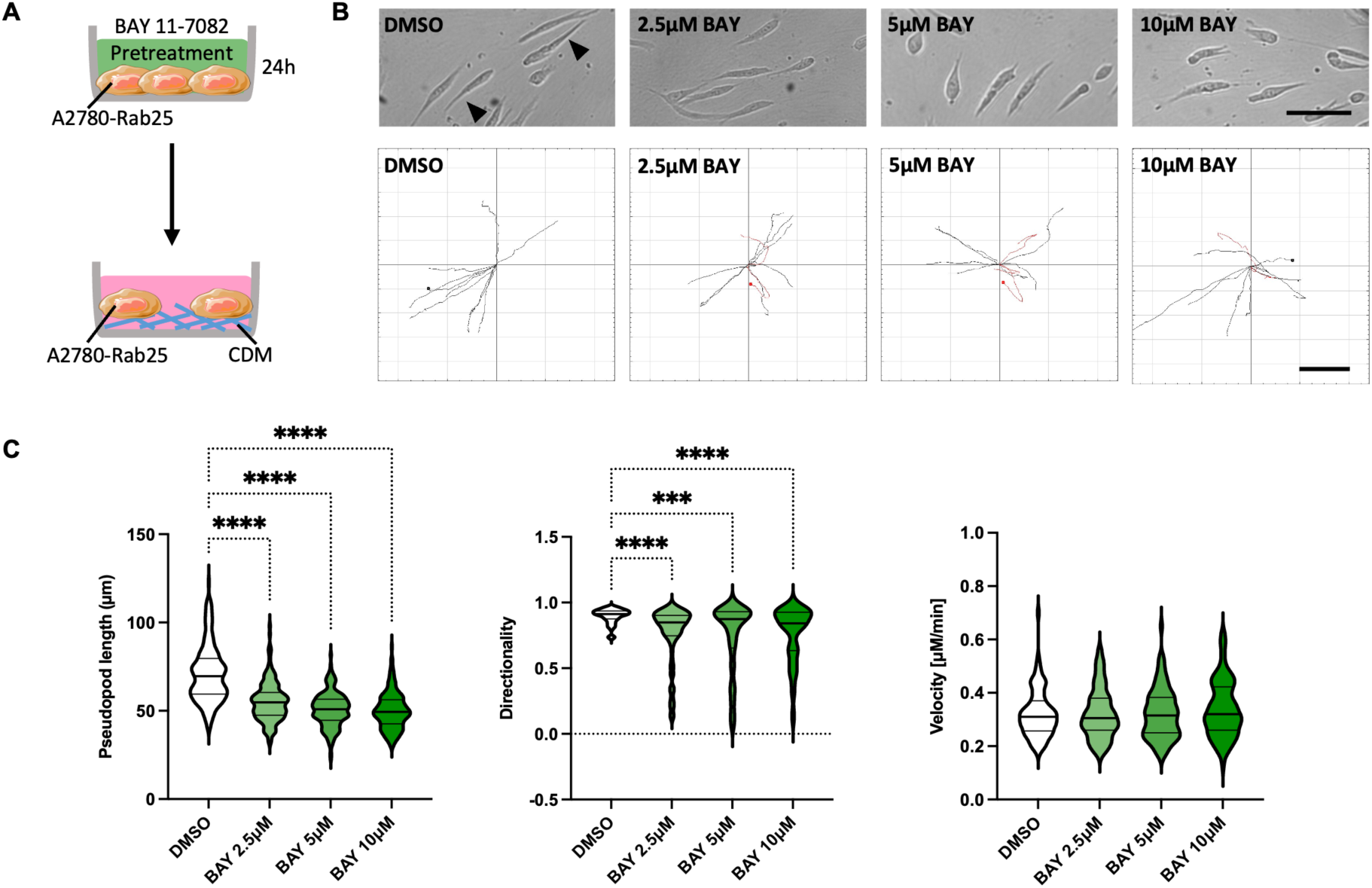
NF-kB inhibition impaired A2780-Rab25 cell migration on CDM. **(A)** Schematic of the experimental workflow. **(B,C)** A2780-Rab25 cells were pretreated with DMSO, 2.5, 5, or 10μM of BAY 11-7072 (BAY) for 24hr, seeded on CDM and imaged live with a Nikon widefield live-cell system (Nikon Ti eclipse with Oko-lab environmental control chamber) for 16h. Stills extracted from the movies are presented. The arrow-heads point to the elongated pseudopods. Scale bar, 100μM. Representative spider plots show the migration paths of manually tracked cells (directionality >0.5 in black, <0.5 in red). Scale bar, 200μM. The pseudopod length (μm), directionality and velocity [μM/min] of cell migration were measured with ImageJ. Data were plotted as violin plots (median and quartiles) from N=2 independent experiments. ***p=0.0002 ****p<0.0001, Kruskal-Wallis test. Image created with items adapted from Servier Medical Art, licensed under CC BY 4.0.

### ADAMTS5 was required for Rab25-dependent 3D invasion of OC cells

As ADAMTS5 was required for Rab25-dependent cell migration, we investigated the role of ADAMTS5 in OC cell invasion in 3D systems. A2780-Rab25 cell spheroids were embedded into a matrix mix containing 3mg/mL Geltrex, 3mg/mL collagen I and 25μg/mL fibronectin in the presence of DMSO, 5 or 10μM ADAMTS5 inhibitor and imaged at day 0, day 1 and day 2 (**Figure 6A**). The area of invading protrusions outside of the spheroid core was marked as the “invasion area” and was normalised to the core area. Treatment with the ADAMTS5 inhibitor resulted in a dose-dependent reduction in cell invasion (**Figure 6B**).

**Figure 6.**
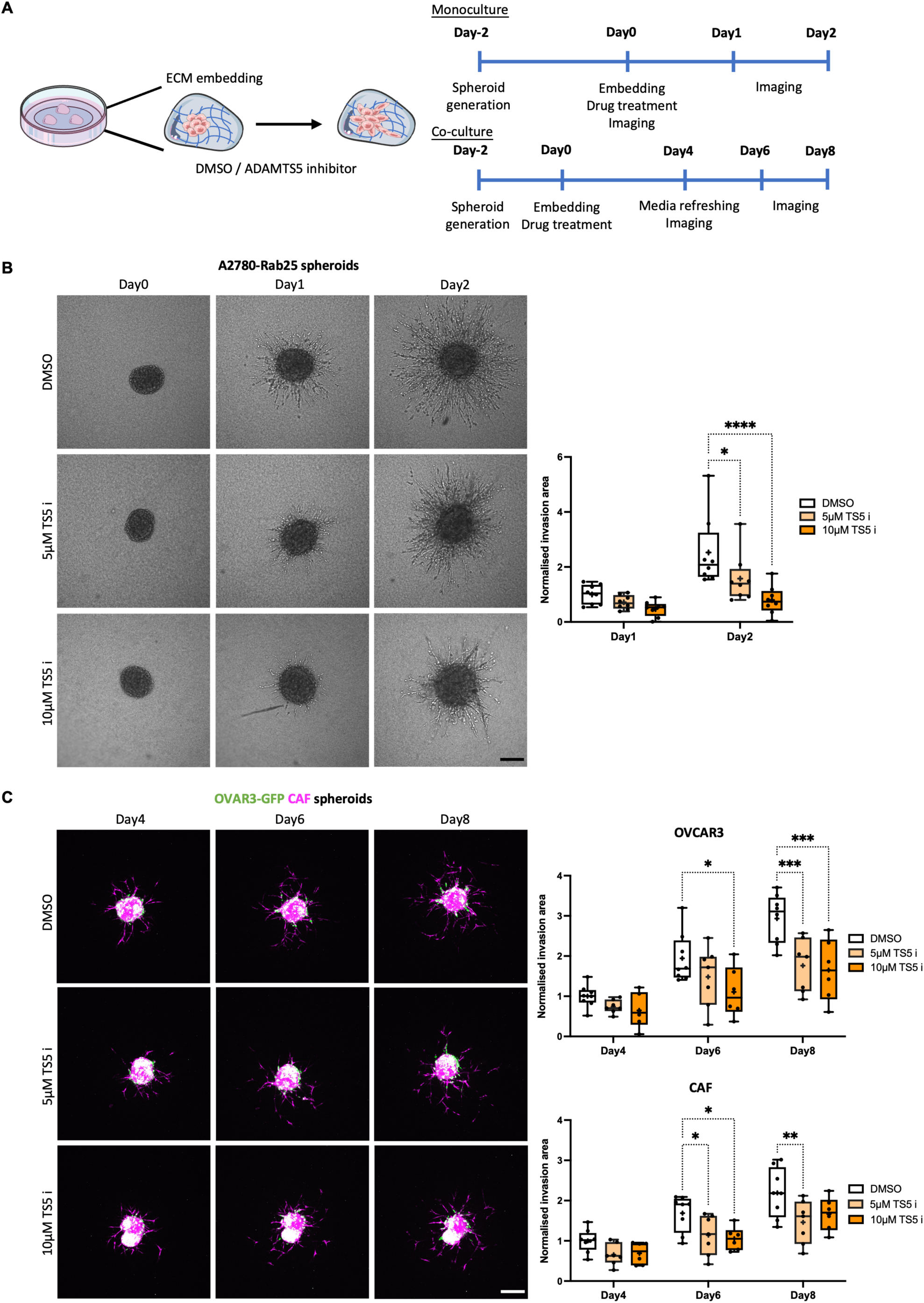
ADAMTS5 inhibition reduced the 3D invasion of OC cells in monoculture and co-culture with CAFs. **(A)** Schematic of 3D monoculture and co-culture invasion assays. (B) A2780-Rab25 spheroids generated by the hanging drop method were embedded in 3mg/mL Geltrex, 3mg/mL collagen I and 25μg/mL fibronectin in the presence of DMSO, 5 or 10|j.M ADAMTS5 inhibitor (TS5 i) and imaged live with a Nikon Al confocal microscope up to day 2. Scale bar, 200μM. Spheroid invasion area was quantified with ImageJ and normalised to DMSO day 1. Data are presented as box and whisker plots (Min to Max, + represents the mean) from N=4 independent experiments. *p=0.0207, ****p<0.0001, two-way ANOVA, Dunnett’s multiple comparisons test. **(C)** OVCAR3 cells stably expressing nuclear GFP (H2B-GFP) and CAFs labelled with Cell tracker™ Red CMTPX (2:1 ratio) were generated, embedded as in B and imaged live with a Nikon Al confocal microscope up to day 8. Scale bar, 200μM. Spheroid invasion area was quantified with ImageJ and normalised to DMSO day 4. Data are presented as box and whisker plots (Min to Max, + represents the mean) from N=3 independent experiments. *p<0.05, **p=0.0035, ***p<0.001, two-way ANOVA, Dunnett’s multiple comparisons test. Image created with items adapted from Servier Medical Art, licensed under CC BY 4.0.

During OC metastasis, CAFs were found to form heterotypic spheroid with tumour cells and promote OC cell invasion through the secretion of proteases and ECM components [25–27]. To better investigate the role of CAFs in ADAMTS5-mediated OC cell invasion, we established a co-culture 3D spheroid model with Cell Tracker Red-labelled CAFs and OVCAR3-GFP cells. The spheroids were embedded into the matrix mix in the presence of DMSO, 5 or 10μM of ADAMTS5 inhibitor and imaged at day 4, day 6 and day 8. Consistent with the A2780-Rab25 cell spheroid results, the invasion of OVCAR3 cells in co-culture with CAFs was impaired by ADAMTS5 inhibition in a dose-dependent manner, indicating that ADAMTS5 catalytic activity was required for Rab25-overexpressing OC cell invasion. Interestingly, CAF invasion was also impaired by ADAMTS5 inhibition in co-culture 3D spheroids (**Figure 6C**).

In our co-culture model, CAFs were observed at the tips of invading strands, which seemed to lead to the invasion of OVCAR3 cells. Since the inhibition of ADAMTS5 reduced the invasion capacity of both CAFs and OVCAR3 cells, it is possible that ADAMTS5 inhibitor indirectly suppressed the invasion of OVCAR3 cells by preventing CAF invasion. To investigate this, monoculture CAF spheroids were generated and embedded in the presence of DMSO or 10μM ADAMTS5 inhibitor. Interestingly, CAF invasion was not affected by ADAMTS5 inhibition in monoculture (**Figure S4**), suggesting that ADAMTS5 might control CAF/OVCAR3 cells crosstalk. Given the timeframe of the 3D spheroid assays, changes in cell proliferation could affect the quantification of the invasion area. To investigate whether ADATMS5 inhibition perturbed cell growth, we performed a 5-ethynyl-2ʹ-deoxyuridine (EdU) incorporation assay in OVCAR3/CAF spheroids (**Figure S5A**). EdU is a thymidine analogue, which is incorporated in the DNA during replication in the S phase of the cell cycle. As a result, only OVCAR3 cells were positive for EdU, and the percentage of EdU positive cells was not affected by ADAMTS5 inhibition (**Figure S5B**), indicating that ADAMTS5 was required for 3D invasion without affecting cell proliferation.

In the TME, both tumour cells and stroma cells including CAFs were found to secrete proteases and remodel the ECM [6]. Therefore, it is possible that ADAMTS5 secreted from both OVCAR3 and CAFs promoted spheroid invasion. To investigate the contribution of ADAMTS5 secreted by cancer cells, OVCAR3-GFP cells were transfected with a non-targeting siRNA control, Rab25 targeting or ADAMTS5 targeting siRNAs and the 3D invasion of co-culture spheroids was measured (**Figure 7A**). While co-culture spheroids generated from control siRNA transfected cells invaded into the surrounding matrix over time, both Rab25 and ADAMTS5 knockdown significantly reduced the invasion ability of OVCAR3 cells in co-culture with CAFs (**Figure 7B**). These data demonstrated that Rab25 and ADAMTS5 expression in cancer cells were required to promote spheroid invasion. Interestingly, Rab25 knockdown in OVCAR3 cells also reduced the invasion ability of CAFs in co-culture, while ADAMTS5 knockdown resulted in a small but not statistically significant inhibition, indicating that Rab25 may be involved in mediating the crosstalk between OC cells and CAFs during metastasis. Altogether, these results show that ADAMTS5 was required for Rab25-dependent invasion of OC cells in 3D systems.

**Figure 7.**
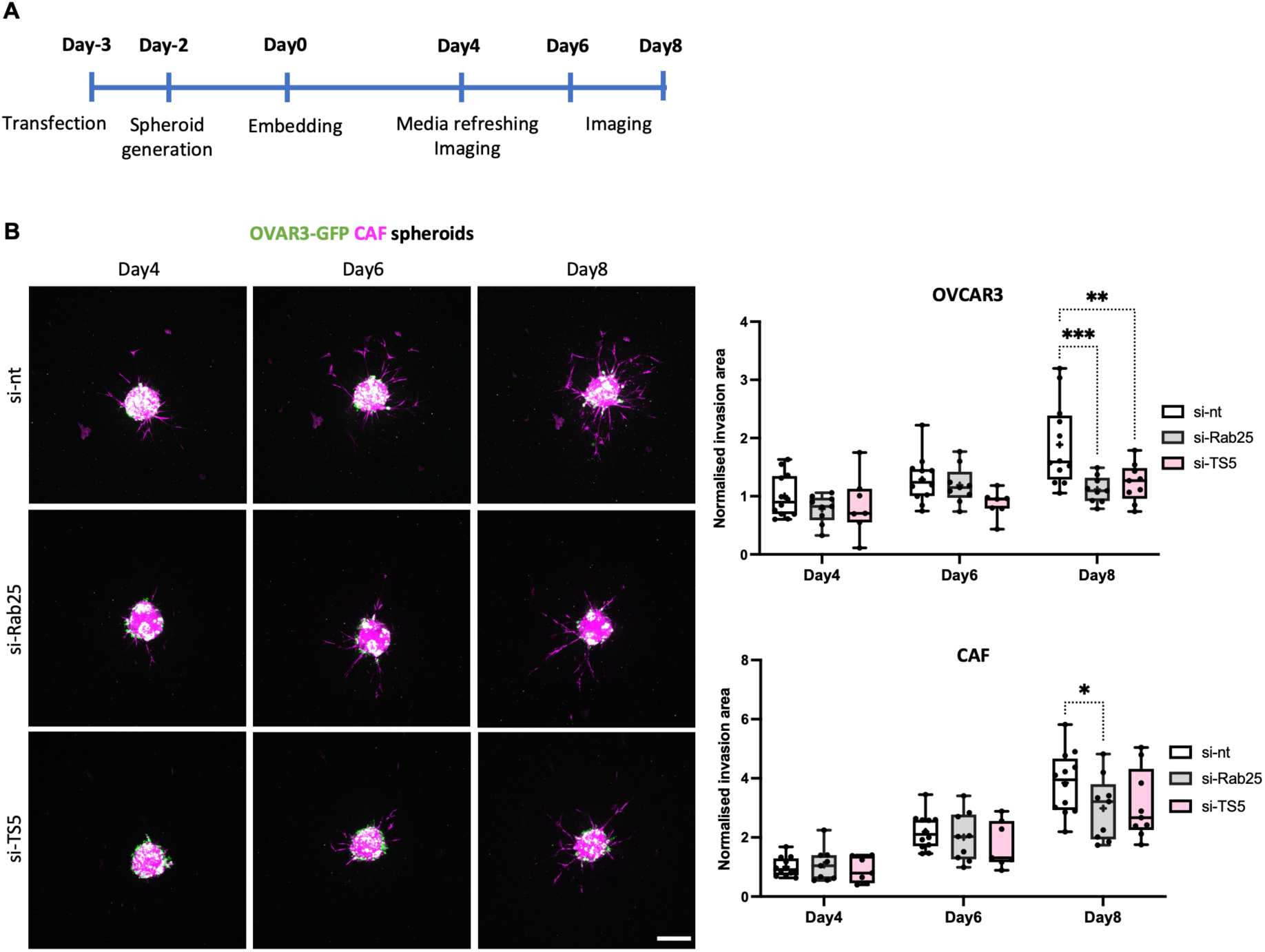
Rab25 and ADAMTS5 KD reduced the 3D invasion of OVCAR3 cells in co-culture. **(A)** Experimental timeline. **(B)** Co­culture spheroids were generated with OVCAR3-GFP cells (green) transfected with a non-targeting control (si-nt), Rab25-targeting (si-Rab25) or ADAMTS5 targeting (si-TS5) si-RNA and Cell tracker™ Red CMTPX (magenta) labelled CAFs (2:1 ratio), embedded in 3mg/mL Geltrex, 3mg/mL collagen I and 25μg/mL fibronectin and imaged live with a Nikon Al confocal microscope up to day 8. Scale bar, 200μM. The spheroid invasion area was calculated with ImageJ and normalised to si-nt day 4. Data are presented as box and whisker plots (Min to Max, + represents the mean) from N=4 independent experiments. *p=0.0353, **p=0.0012, ***p=0.0001, two-way ANOVA, Dunnett’s multiple comparisons test.

### ADAMTS5 is a poor prognosis factor for ovarian cancer

Given the role of ADAMTS5 in the migration and invasion of OC cells, we looked at the relationship between ADAMTS5 expression and survival outcomes of OC patients. Interestingly, high ADAMTS5 expression correlated with reduced overall survival (OS), progression-free survival (PFS) and the palliative performance scale (PPS) (**Figure 8A-C**). Taken together, these data suggest that ADAMTS5 could promote metastasis by enhancing OC cell migration and invasion, leading to poor prognosis outcomes for OC patients. Therefore, ADAMTS5 might represent a novel therapeutic target to prevent of OC metastasis.

**Figure 8.**
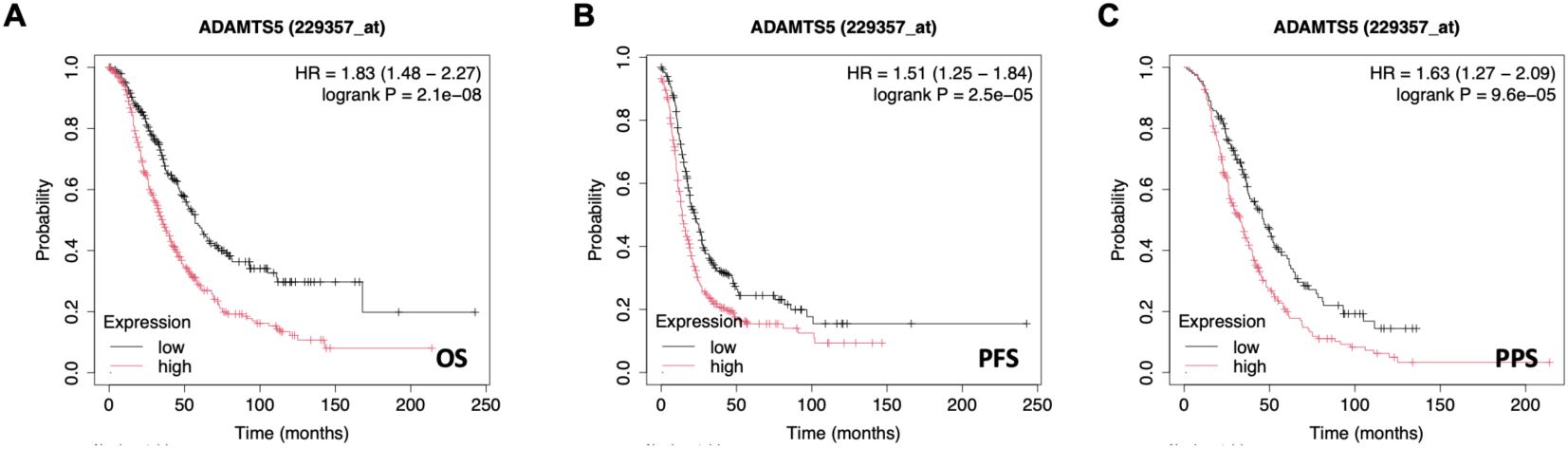
ADAMTS5 expression correlated with with poor prognosis of OC patients. **(A-C)** 655 ovarian cancer patients were stratified into low and high ADAMTS5 expression. The Kaplan-Meier analysis compared **(A)** overall survival (OS), **(B)** progression­free survival (PFS) and **(C)** palliative performance scale of OC patients with tumours expressing high ADAMTS5 levels (red) with those expressing low (black) ADAMTS5 levels.

## Discussion

Altered ECM composition and structure have been widely detected in the TME of multiple solid tumours, including ovarian cancer [26]. These changes are mainly caused by the dysregulation of ECM-modifying enzymes, which can be secreted by both tumour cells and CAFs [6, 28]. Here, we demonstrated that Rab25 induced ADAMTS5 expression in OC cells, by upregulating the activity of NF-κB, in an AKT- and ERK-independent manner. Remarkably, we showed that ADAMTS5 pharmacological inhibition or knockdown significantly reduced OC cell migration and invasion, without affecting cell proliferation (**Figure 9**).

**Figure 9.**
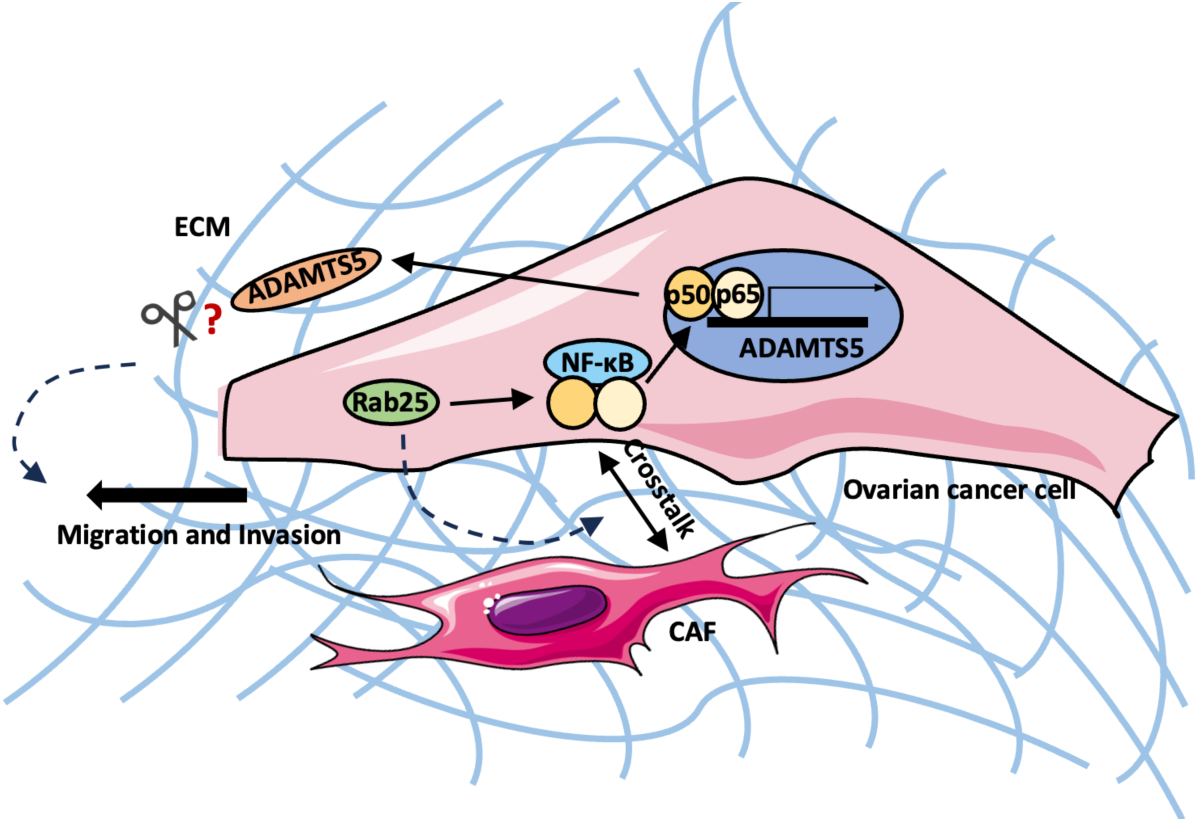
Working model. Upregulated Rab25 induced ADAMTS5 expression through an NF-KB-depended signalling pathway in OC cells. Secreted ADAMTS5 promoted OC cell migration and invasion, in a proteolytic activity-related manner. Rab25 could also be involved in controlling the crosstalk between OC cells and CAFs in the TME. Image created with items adapted from Servier Medical Art, licensed under CC BY 4.0.

We found that both endogenous and exogenous overexpression of Rab25 increased ADAMTS5 expression, while the presence of the ECM further increased ADAMTS5 levels in A2780 but not OVCAR3 cells.

This could be due to the different origins of the cells, as A2780 were established from an ovarian endometrioid adenocarcinoma sample, while OCVAR3 were derived from the ascitic fluid of an OC patient with high-grade ovarian cancer [29]. Unfortunately, no reliable antibody is currently available to study the protein levels of ADAMTS5 (S. Santamaria, personal communication), therefore we were unable to test ADAMTS5 secretion in the conditioned media of OVCAR3 cells.

In agreement with data in chondrogenic cells, here we showed that NF-κB was required for ADAMTS5 expression in OC cells overexpressing Rab25. While a direct interaction between NF-κB and the ADAMTS5 promoter has been reported [17], we cannot rule out the possibility of an indirect regulation of ADAMTS5 expression, via the modulation of cytokines, such as IL-1, IL-6 and IL-8 downstream of NF-κB activation [30].

Indeed, IL-1β and IL-6 were shown to induce ADAMTS5 expression in osteoarthritis [19]. Although NF-κB has been shown to be regulated by PI3K/AKT and MAPK/ERK pathways in OC cells [22, 31], none of these were dependent on Rab25 in our settings. This indicates that Rab25 might activate NF-κB via other signalling pathways, which warrant further investigation. It has been suggested that the mammalian target of rapamycin complex 1 (mTORC1) might regulate NF-κB. Interestingly, we have demonstrated that Rab25 promotes mTORC1 activation in OC cells [24], suggesting that Rab25 might control NF-κB in a mTORC1-dependent manner.

We found that the catalytic activity ADAMTS5 was required for cell migration and invasion, in the presence of ECM, as we could inhibit both processes using a pharmacological inhibitor that blocks ADAMTS5 ability to cleave its substrates. ADAMTS5 was found to cleave aggrecan, brevican, neurocan and versican [32]. Among them, versican was found to be increased in OC stroma, which was shown to stimulate OC cell migration and invasion and promote peritoneal metastasis [9, 33, 34]. In addition, versican secreted by primary ovarian CAFs was found to promote OC cell invasion [35]. Since blocking the ADAMTS cleavage site on versican reduced the migration of glioma cancer cells [36], it is possible that ADAMTS5 could also drive OC cell migration by cleaving its substrate versican. Interestingly, versican cleavage by ADAMTSs results in the release of a matrikine fragment, versikine, which has been shown to promote glioma cell migration [36]. Further work will elucidate the contribution of versican cleavage to OC cell migration.

Interestingly, we found that NF-κB was required for Rab25-dependent cell migration on CDM. This is consistent with previous findings, highlighting a pro-migratory role of NF-κB in lung, breast and ovarian cancer cells [37–40]. In addition, NF-κB inhibition also significantly impaired metastasis formation in mouse xenograft models [38].

Additionally, our data showed that the knockdown of Rab25, and to a lesser extent ADAMTS5, in OC cells reduced the CAF invasion in co-culture spheroids, while ADAMTS5 inhibition did not affect CAF invasion in mono-culture spheroids, suggesting a potential role of Rab25 and ADAMTS5 in controlling cancer cell/CAF crosstalk. To note, Rab25 is epithelial-specific and, therefore, not expressed in CAFs [41]. Transforming growth factor β (TGFβ) has a key role in sustaining CAF phenotypes. Interestingly, members of the ADAMTS family, including ADAMTS1, 6 and 10 were found to control TGFβ signalling [42, 43]. Moreover, TFGβ1 expression has been reported to be upregulated in Rab25-expressing cells [10], suggesting that ADAMTS5 and Rab25 might control OC cell/CAF crosstalk via TGFβ signalling.

ADAMTS5 has been reported to either promote or inhibit tumour formation, in a context-dependent manner. Consistent with our analysis showing a correlation between high ADAMTS5 expression and poor patient prognosis in OC, ADAMTS5 was shown to be a tumour promoter in glioblastoma [44], non-small cell lung cancer [45] and head and neck cancer [46], while in melanoma and gastric carcinoma, ADAMTS5 was found to suppress tumour progression by inhibiting angiogenesis, in a catalytic activity-independent manner [47, 48].

In conclusion, our study demonstrated that Rab25 induced ADAMTS5 expression in an NF-κB-dependent manner in OC cells. ADAMTS5 was required for OC cell migration and invasion, and its expression correlated with reduced OC patient survival, suggesting that it might be a novel regulator of OC metastasis. Since small molecule inhibitors of ADAMTS5 are in phase III clinical trials for osteoarthritis treatment [49], our data suggest that these could be repurposed to prevent OC progression.

## Methods

### Reagents

Primary antibodies for western blotting: ADAMTS5 (abcam, ab41037), Rab25 (Proteintech, 18139-1-AP), AKT (Cell Signalling, #4691), p-AKT (Cell Signalling, #4060), ERK1/2 (Cell Signalling, #9102), p-ERK (Cell Signalling, #9101), GAPDH (Santa Cruz Biotechnology, SC-47724) and α-Tubulin (Sigma-Aldrich, T9026). Secondary antibodies for western blotting: IR Dye 680LT anti-Rabbit antibody (LICOR Biosciences), IR Dye 800LT anti-Mouse antibody (LICOR Biosciences). Primary and secondary antibodies for immunofluorescence: NF-κB/p65 (Proteintech,10745-1-AP), Alexa Fluor® 488 goat anti-rabbit IgG (H+L) (Invitrogen, A11034), Alexa Fluor® 555 donkey anti-rabbit IgG (H+L) (Life technologies, A31572). Primers for RT-qPCR: ADAMTS5 (QIAGEN, Hs_ADAMTS5_1_SG), Rab25 (QIAGEN, Hs_RAB25_1_SG), GAPDH (QIAGEN, Hs_GAPDH_1_SG), CXCL8 (QIAGEN, Hs_CXCL8_1_SG). ON-TARGETplus SMARTpool siRNAs (Dharmacon, Horizon discovery): ADAMTS5 (L-005775-00-0005), Rab25 (L-010366-00-0005), siGENOME non-targeting control siRNA #4.

### Cell culture

The OC cell lines A2780, OVCAR3, OVCAR4 and omental cancer-associated fibroblast (CAFs) were cultured in RPMI-1640 medium supplemented with 10% (v/v) foetal bovine serum (FBS) and 1% (v/v) penicillin/streptomycin (Pen/Strep). SKOV3 and telomerase-immortalised human dermal fibroblasts (TIFs) were cultured in high glucose Dulbecco’s modified Eagle medium (DMEM) supplemented with 10% (v/v) FBS and 1% (v/v) Pen/Strep. A2780-DNA3 and Rab25 cells generated in Dr Gordon Mill’s lab as described in [12] were a gift from Prof Jim Norman’s lab (Cancer Research UK Scotland Institute). To maintain the overexpression of Rab25, cell lines were selected with 0.4mg/mL G418 for a week every 10 passages. OVCAR3 cells were a gift from Prof Patrick Caswell’s lab (The University of Manchester) and were STR profiled. The hTERT immortalised omental CAFs from Prof Sara Zanivan’s lab (Cancer Research UK Scotland Institute, Glasgow) were generated as described in [15]. All cell lines were maintained at 37°C in 5% CO_2_, passaged every 3 to 4 days, and routinely tested for mycoplasma contamination.

### Cell-derived matrix (CDM) generation

CDM generation has been previously described [50]. Briefly, plates were coated with 0.2% (v/v) gelatin in PBS for 1 h at 37°C, followed by crosslinking with 1% (v/v) sterile glutaraldehyde in PBS for 30 min at room temperature (RT). The glutaraldehyde was quenched with 1M sterile glycine in dH_2_O for 20 min at RT and complete medium was used for equilibration for 30 min at 37°C. TIFs and CAFs were then seeded on top of the coated plates. After reaching confluency, the media was replaced with complete media containing 50μg/mL ascorbic acid and was refreshed every other day. TIFs were kept in DMEM complete media with ascorbic acid for 9 days to secrete CDM. To extract TIF-CDM, triton extraction buffer (20mM NH_4_OH and 0.5% (v/v) Triton X-100 in PBS containing Ca^2+^ and Mg^2+^ (PBS^++^)) was added on top and left at RT until all cells were removed. Then, 10μg/mL DNase I in PBS^++^ was added and the plates were incubated at 37°C for 1 h. CAFs were kept in Human Plasma-Like Medium (HPLM, Gibco) with ascorbic acid for 11 days. The CDM was extracted with PLA2 extraction buffer (50mM Tris-HCl pH 8, 150mM NaCl, 1mM MgCl_2_, 1mM CaCl_2_, 0.5% sodium deoxycholate and 20 unit/ml PLA2) at 37°C for 1 h. Then, 10μg/mL DNase I in PBS^++^ was added and the plates were incubated at 37°C overnight. The CDMs were stored at 4°C in PBS^++^ and were used within two weeks.

### Conditioned media (CM) harvest

A2780 cells were seeded on plastic and incubated at 37°C in 5% CO_2_ for 3 days. Then, the conditioned media (CM), was spun at 300g for 10 min, 2,000g for 10 min, and 10,000g for 30 min at 4°C. The supernatant was transferred into new falcon tubes and stored at 4°C for up to a week.

### Western blotting

Cells seeded on plastic were lysed in SDS-lysis buffer (50mM Tris PH7, 1% SDS in dH_2_O) and cells seeded on CDM were lysed with triton extraction buffer (20mM NH_4_OH and 0.5% (v/v) Triton X-100 in PBS containing Ca^2+^ and Mg^2+^ (PBS^++^)). Cell lysates were then spun down through QiaShredder columns at full speed for 5 min. To analyse CM proteins, A2780-DNA3 and Rab25 cells were seeded in complete media and cultured at 37°C in 5% CO_2_, and the culture media was changed into serum-free media ∼4hr after seeding. After 3 days, the CMs were subjected to the 3-step centrifugation protocol as described above. The CMs were further concentrated using Amicon Ultra® - 4 Centrifugal filters (3,000 or 10,000 MWCO PES). For both cell lysate and concentrated CM, the samples were mixed with 4x NuPAGE buffer containing 1mM DTT and boiled at 70°C for 5 min. 25μL of the samples and 1μL of the protein ladder (BioLabs) were loaded into a Bio-Red 4-15% Mini-PROTEAN precast polyacrylamide gel, then run at 100V for 75 min in 1x running buffer (25mM Tris, 192mM glycine and 1% SDS in dH_2_O). The membrane transfer was performed in Towbin transfer buffer (25mM Tris, 192mM glycine and 20% methanol (v/v) in dH_2_O, pH 8.3) at 100V for 75 min. The membranes were then blocked with 5% (w/v) milk in 1xTBST (50mM Tris HCl, 150mM NaCl and 0.5% (w/v) Tween 20 in dH_2_O) for 1 h at RT. The membranes were incubated overnight with primary antibodies (ADAMTS5 1:250, Rab25 1:600, AKT 1:1000, p-AKT 1:1000, ERK 1:1000 and p-ERK 1:1000) together with GAPDH (1:1000) in 1xTBST at 4°C, followed by 1-h incubation at RT with secondary anti-mouse IgG LICOR IR Dye 800 (1:30,000) and anti-rabbit IgG LICOR IR Dye 680 (1:20,000) in TBST with 0.01% (w/v) SDS. The Membranes were then imaged with a LICOR Odyssey Sa system, and the intensity of the protein bands was quantified with Image Studio Lite software.

### RT-qPCR

mRNA was extracted from snap-frozen cell pellets according to the manufacturer’s protocol (RNeasy® Mini – QIAGEN). cDNA was then synthesised using the High-Capacity cDNA Reverse Transcription Kit (Fisher) following the manufacturer’s protocol. Then, a final concentration of 1x QuantiNova SYBR® Green PCR Kit (QIAGEN) master mix and 1x QuantiTect® Primer Assay for target genes (ADAMTS5, Rab25 and GAPDH) was prepped in RNase-free water. 7μL of loading master mix and 3μL of cDNA solution (1:100 dilution, 5ng/μL final conc.) were loaded into a 384-well plate. The -RT and blank water controls were tested for all target genes in each individual running. Quantstudio 12K flex real-time PCR system was used to analyse the samples in the SYBR® mode. Expression levels of the target genes were calculated using 2^-ΔΔCt^ method (2^-(ΔCt Target gene – ΔCt Housekeeping gene)^) with GAPDH as the housekeeping gene. Each sample was tested in three technical replicates.

### siRNA transfection

2μL Dharmafect I (Dharmacon) was mixed with 198μL FBS & antibiotic-free RPMI medium and incubated for 5 min at RT. Meanwhile, 2.5μL of 20μM siRNA and 197.5μL FBS & antibiotic-free RPMI medium were mixed, added to the Dharmafect solution and incubated for 20 min at RT. The 400μL solution was then added on top of the cells and topped up to 2mL complete medium without Pen/Strep. 24 h after transfection, the media was changed to complete medium without Pen/Strep. siGENOME non-targeting control siRNA #4 was used as a non-targeting control.

### Immunofluorescence

35mm glass-bottom dishes (12mm glass diameter, NEST®) were coated with 0.1mg/mL Geltrex (Gibco) to facilitate cell adhesion. Cells were fixed with 4% (v/v) paraformaldehyde (PFA, Thermo Scientific) in PBS for 15 min and permeabilised with 0.25% (v/v) Triton X-100 in PBS for 5 min at RT. Cells were blocked with 1% (w/v) BSA in PBS for 1 h, stained with primary anti-NF-κB/p65 antibody (1:100 in 1% BSA/PBS) for 1h at RT, and incubated with secondary antibody (Alexa Fluor® 488 goat anti-rabbit IgG (H+L) or Alexa Fluor® 555 donkey anti-rabbit IgG (H+L), 1:1000 in 1% BSA/PBS) for 45 min. Since GFP-OVCAR3 cells were used in these experiments, the Alexa Fluor 488-conjugated secondary antibody was used for A2780 cells, while the Alexa Fluor 555-conjugate one was used for OVCAR3 cells. For simplicity, NF-κB/p65 staining is presented in green for both cell lines in the result section. The actin cytoskeleton was stained with Phalloidin Alexa Fluor 555 (1:400 in PBS) or Alexa Fluor 647 (1:300 in PBS) for 10 minutes. Finally, Vectashield mounting agent containing DAPI was added on top. Images were taken with a Nikon A1 confocal microscope, CFI Plan Apochromat VX 60X oil immersion objective and the integrated density was measured in Fiji/Image J [51].

### Cell migration assay

Cells were seeded on plastic or 12-well plates containing CDM and allowed to spread for 4 h before imaging. For the CM migration assays, the cells were seeded in complete media mixed with CM (1:1 dilution). Where indicated, DMSO or inhibitors were added into the culture media before imaging. Cells were imaged live with a Nikon widefield live-cell system (Nikon Ti eclipse with Oko-lab environmental control chamber) with a Plan Apo 10X objective (NA 0.75). Images were acquired every 10 min for 16 h. For each condition, 5 positions were imaged randomly. Cell migration was manually tracked with ImageJ plugin Manual Tracking [51, 52], and the migration velocity and directionality were calculated with the chemotaxis tool plugin (https://ibidi.com/chemotaxis-analysis/171-chemotaxis-and-migration-tool.html). Pseudopod length, which describes the distance between nuclei and the tip of the pseudopod in the direction of cell migration, was measured with the “Straight” tool in ImageJ.

### 3D spheroid invasion assay

Spheroids were generated using the hanging drop method as previously described [53]. Briefly, A2780-Rab25 cells and CAFs were labelled with Cell tracker^TM^ Red CMTPX (1:3000 in FBS-free media) at 37°C for 1 h. OVCAR3 cells were transfected with pCAG-H2B-GFP plasmid (Addgene #184777) using Invitrogen^TM^ Lipofectamine^TM^ 2000 Transfection Reagent following the manufacturer’s protocol and a stable cell line was established. Then, 1×10^5^ (monoculture) or 2.1×10^5^ (co-culture, OVCAR3:CAF=2:1) labelled cells were suspended in 2mL of medium containing 20μg/mL of soluble collagen I (BioEngineering) and 4.8mg/mL of Methyl Cellulose (MTC, Sigma-Aldrich). 20μL drops of the cell suspension were then hanged on the lid of a 10cm tissue culture dish to generate the spheroids. After 48 h at 37°C, the spheroids were harvested, embedded into 45μL of a matrix mix containing 3mg/mL collagen I (Ibidi), 3mg/mL Geltrex (Gibco) and 25μg/mL Fibronectin (Sigma-Aldrich), and placed on a 35mm glass-bottom dish. During matrix polymerisation, the dishes were incubated at 37°C upright for 2 min, then flipped upside down and incubated for another 2 min. After 5 up and down flips (10 minutes in total), the dishes were kept upside down and incubated at 37°C for 20 to 25 min until the matrices polymerised. Spheroids were cultured in complete media containing DMSO, 5 or 10μM ADAMTS5 inhibitor. Spheroids were imaged live with Nikon A1 confocal microscope, CFI Plan Fluor 10x objective (NA 0.3). The fluorescent images were thresholded in ImageJ and the areas of the spheroid cores and the total area were measured. The invasion area was then calculated by total area – core area, and the invasion area was normalised to the core area.

### 3D EdU incorporation assay

Co-culture spheroids were generated with OVCAR3-GFP cells and unlabelled CAFs, embedded and treated with DMSO or ADAMTS5 inhibitor as described above. After 6 days, a final concentration of 10μM EdU solution was added to the media. After 2 days, the spheroids were fixed with 4% PFA containing Hoechst 33342 (1:500) at 37°C for 20 min. Then, the spheroids were permeabilised with IF wash buffer (46mM NaN_3_, 0.1% (w/v) BSA, 0.2% (v/v) Triton-X 100 and 0.04% (v/v) Tween-20 in PBS) for 2 h at RT. The spheroids were then stained with Click-iT EdU Imaging Kits (Invitrogen) at 4°C overnight. The spheroids were washed twice with PBS and imaged with a Nikon A1 confocal microscope, CFI Plan Fluor 10x objective (NA 0.3) and the thresholded area of Hoechst, GFP and EdU signal were measured in ImageJ.

### Survival analysis

The survival analysis was performed with Kaplan-Meier plotter (https://kmplot.com/analysis/) [54], using ovarian cancer mRNA gene-chip data. Sources for the databases include Gene Expression Omnibus (GEO), European Genome-Phenome Archive (EGA), and The Cancer Genome Atlas (TCGA).

### Statistical analysis

Graphs were generated with GraphPad Prism software (version 9.1.0). Bar graphs are presented as super-plots [55], where single data points from individual experiments are presented in different shades of blue or orange and the means of each biological replicate are show as black dots. To compare 2 datasets, Mann-Whitney test was used; to compare more than 2 datasets, one-way ANOVA (Kruskal-Wallis test) was used when there was one independent variable, while 2-way ANOVA (Dunnett’s test) was performed when there were 2 independent variables.

## Acknowledgements

We thank Prof Caswell for gifting the OVCAR3 cells, Prof Zanivan for gifting the hTERT immortalized omental CAFs and Prof Barbaric for gifting the pCAG-H2B-GFP plasmid. Imaging work was performed at the Wolfson Light Microscopy Facility, University of Sheffield, using the Nikon A1 confocal and Nikon widefield microscope. qPCR analysis was performed in collaboration with the Tsakiridis lab at the University of Sheffield. We thank Dr Matthews, Dr Maib and Prof King for advice and feedback provided during lab meetings. E.R. is funded by CRUK (C52879/A29144). S.Y. is funded by University of Sheffield Faculty of Science Top Up Scholarship. The Wolfson Light Microscopy Facility, University of Sheffield, is funded by the Wellcome Trust (grant WT093134AIA).

## Supplementary figures

**Figure S1.**
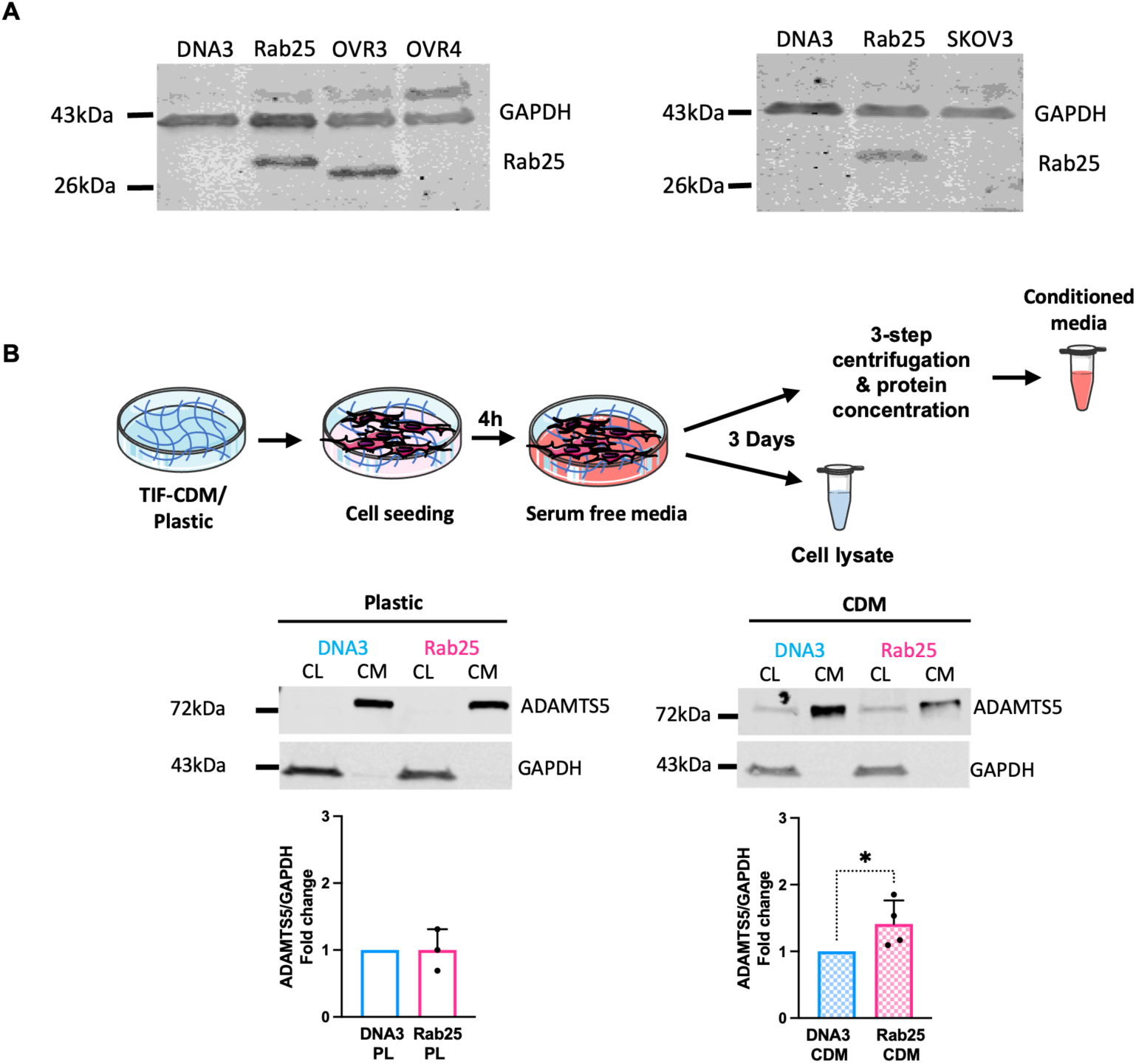
Rab25 induced ADAMTS5 protein expression in OC cells. **(A)** A2780-DNA3, A2780-Rab25, OVCAR3, OVCAR4 and SKOV3 cells were seeded on plastic,Rab25 and GAPDH protein levels were quantified by Western Blotting. Membranes were imaged with a Licor Odyssey Sa system. N=1 independent experiment. (B) Schematic, cell lysate (CL) and conditioned media (CM) harvesting. **(C)** A2780-DNA3 and A2780-Rab25 cells were plated on plastic (PL, N=3 independent experiments) or TIF-CDM (CDM, N=4 independent experiments), cell lysates and concentrated conditioned media were extracted, protein levels of ADAMTS5 in CL and CM, and GAPDH in CL was measured by Western Blotting. Membranes were imaged with a Licor Odyssey Sa system, and the band intensity was quantified by Image Studio Lite software. Data are presented as mean ± SEM. *p=0.0286, Mann-Whitney test. Image created with items adapted from Servier Medical Art, licensed under CC BY 4.0.

**Figure S2.**
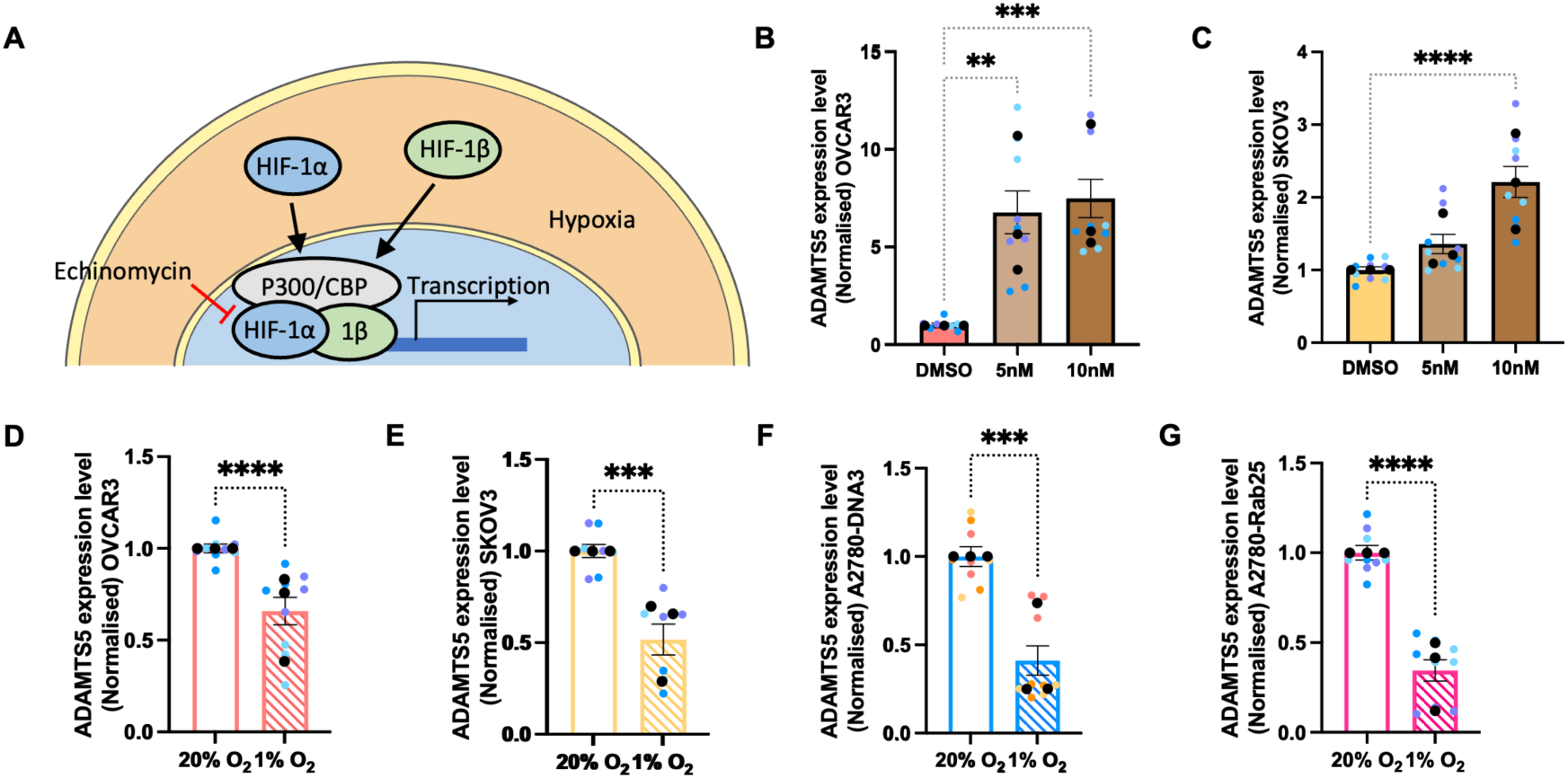
HIF-1 suppressed ADAMTSS expression in OC cells. **(A)** Schematic, HIF-1 signalling pathway. **(B,C)** OVCAR3 **(B)** and SKOV3 (C) cells were treated with DMSO, 5 or 10μM Echinomycin for 24hr and the mRNA levels of ADAMTSS were measured by qPCR. Data were normalised to the DMSO control and presented as mean ± SEM from N=3 independent experiments. The black dots represent the mean of individual experiments. **p=0.0011, ***p=0.0003, ****p<0.0001, Kruskal-Wallis test. **(D-G)** OVCAR3 **(D),** SKOV3 **(E),** A2780-DNA3 **(F)** and A2780-Rab25 **(G)** cells were incubated under Normoxia (20% 0_2_) or Hypoxia (1% 0_2_) for 24hr and the mRNA levels of ADAMTSS were measured by qPCR. Data were normalised to Normoxia control and presented as mean ± SEM from N=3 independent experiments. The black dots represent the mean of individual experiments. ***p<0.001, ****p<0.0001, Mann-Whitney test.

**Figure S3.**
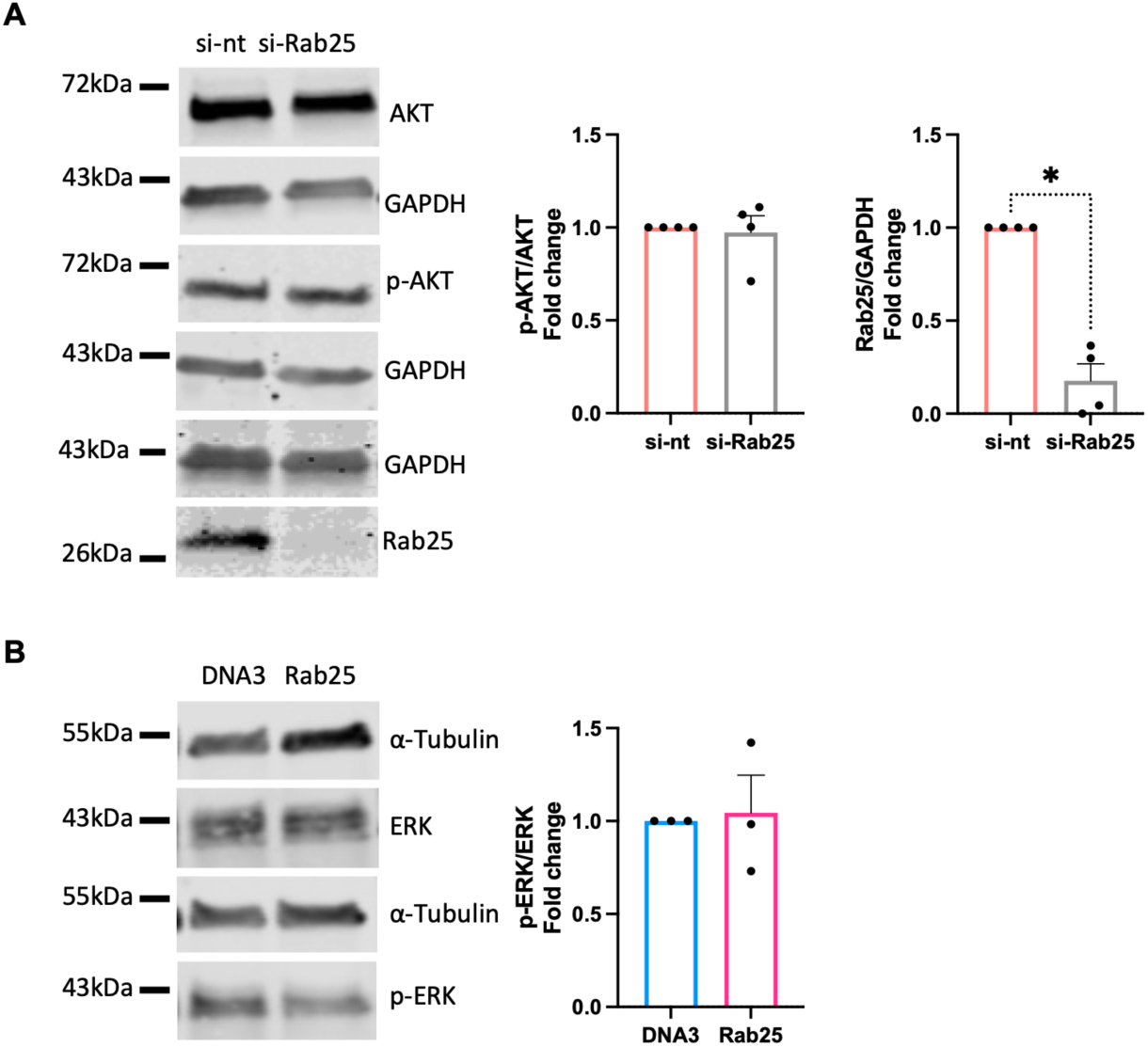
Rab25 downregulation did not affect AKT phosphorylation. **(A)** OVCAR3 cells were transfected with a non-targeting siRNA control (si-nt) or Rab25 targeting siRNA (si-Rab25) and the protein levels of AKT, p-AKT, Rab25 and GAPDH were quantified by Western Blotting. Membranes were imaged with a Licor Odyssey Sa system, and the band intensity was quantified by Image Studio Lite software. The fold change of normalised p-AKT/AKT and Rab25/GAPDH was plotted. Data are presented as mean ± SEM from N=4 independent experiments. *p=0.0286, Mann-Whitney test. **(B)** The protein levels of ERK, p-ERK and a-Tubulin in A2780-DNA3 and A2780-Rab25 cells were quantified by Western blotting as in A. The fold change of normalised p-ERK/ERK was plotted. Data are presented as mean ± SEM from N=3 independent experiments. Mann-Whitney test.

**Figure S4.**
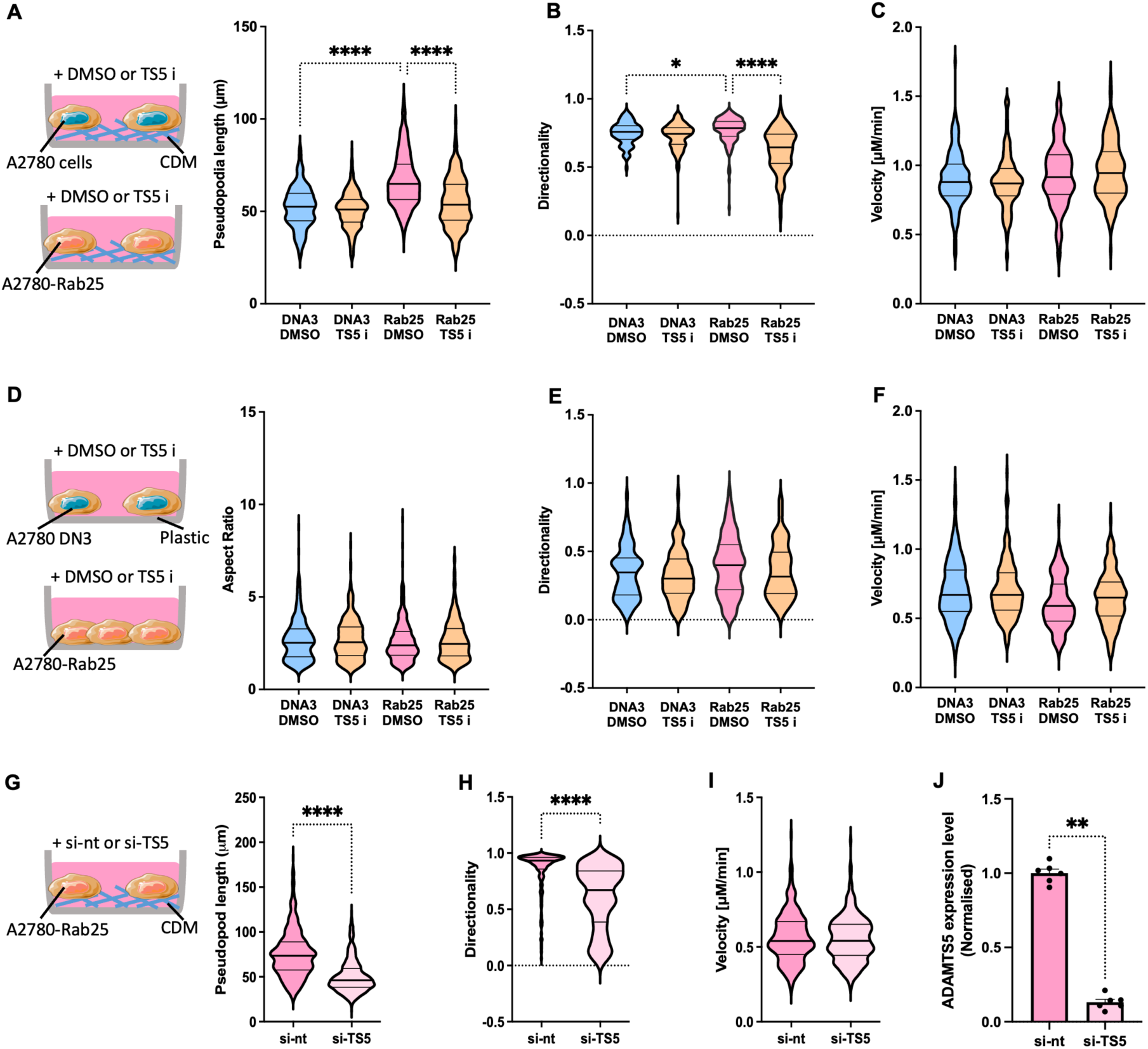
ADAMTSS was required for Rab25-dependent pseudopod elongation and directional migration of OC cells on CDM. **(A-C)** A2780-DNA3 and A2780-Rab25 cells were seeded on CDM and treated with DMSO or SµM ADAMTSS inhibitor (TSS i). Cells were imaged live with a 10X Nikon widefield live-cell system (Nikon Ti eclipse with Oko-lab environmental control chamber) for 16hr. The pseudopod length (µm) **(A),** directionality **(B)** and velocity [µM/min] (C) were measured with lmageJ. Data were plotted as violin plots (median and quartiles) from N=3 independent experiments. *p=0.0155, ****p<0.0001, Kruskal-Wallis test. **(D-F)** A2780-DNA3 and A2780-Rab25 cells were seeded on plastic, treated and imaged as in A. The aspect ratio, measured as cell length/cell width **(D),** directionality **(E)** and velocity [µM/min] **(F)** were measured with lmageJ. Data were plotted as violin plots (median and quartiles) from N=2 independent experiments. **(G-1)** A2780-Rab25 cells were transfected with non-targeting siRNA control (si-nt) or ADAMTSS targeting si-RNA (si-TSS) and seeded on CDM. Cells were imaged as in A and the pseudopod length (µm) **(D),** directionality **(E)** and velocity [µM/min] **(F)** were measured with lmageJ. Data were plotted as violin plots (median and quartiles) from N=3 independent experiments. ****p<0.0001, Mann-Whitney test. **J.** A2780-Rab25 cells were transfected as in D and the mRNA levels of ADAMTSS and GAPDH were measured by qPCR. Data were normalised to si-nt and presented as mean ± SEM from N=2 independent experiments. **p=0.0022, Mann-Whitney test. Image created with items adapted from Servier Medical Art, licensed under CC BY 4.0.

**Figure S5.**
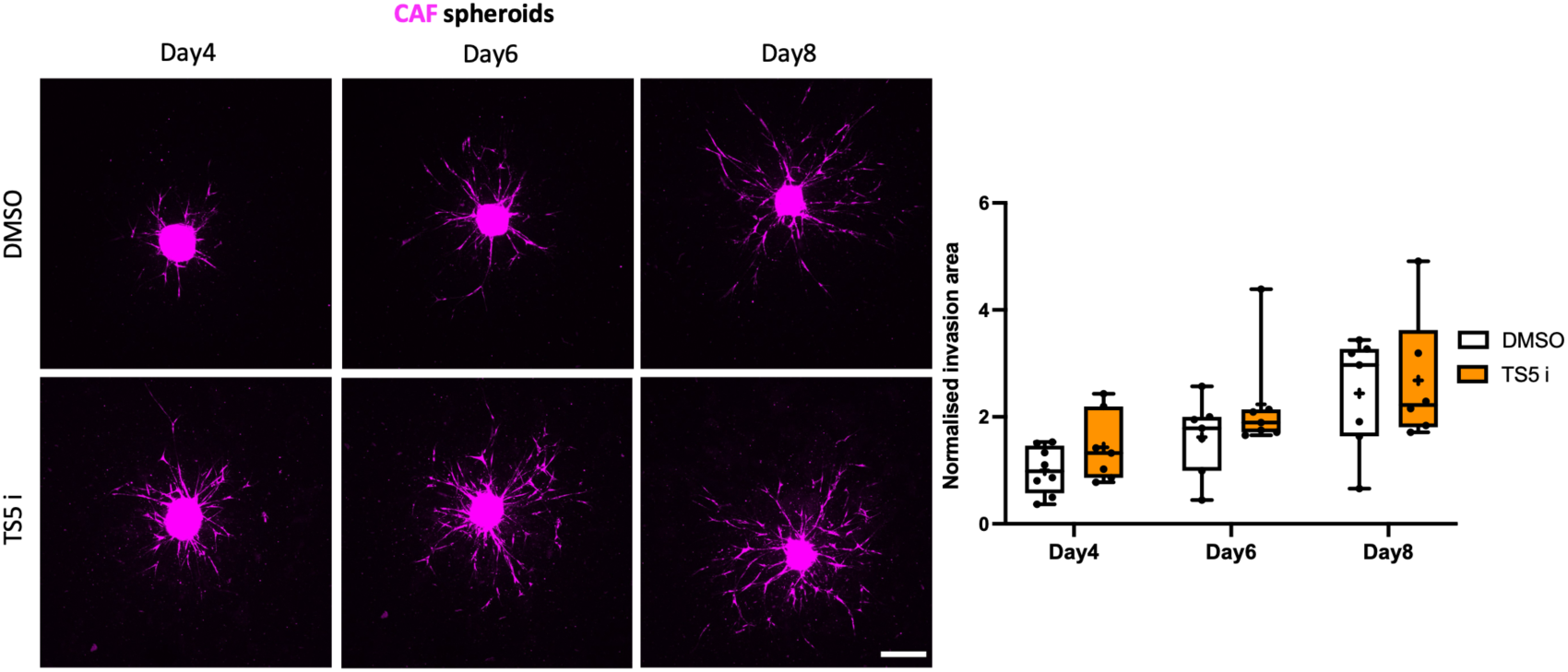
ADAMTSS inhibition did not affect CAF spheroid 3D invasion. Spheroids generated with Cell tracker™ Red CMTPX labelled CAFs were embedded in 3mg/ml Geltrex, 3mg/mL collagen I and 25µg/ml fibronectin in the presence of DMSO control or 10µM ADAMTSS inhibitor (TSS i) and imaged live with a Nikon Al confocal microscope up to day 8. Scale bar, 200µM. Spheroid invasion area was quantified with lmageJ and normalised to DMSO day 4. Data are presented as box and whisker plots (Min to Max, + represents the mean) from N=3 independent experiments.

**Figure S6.**
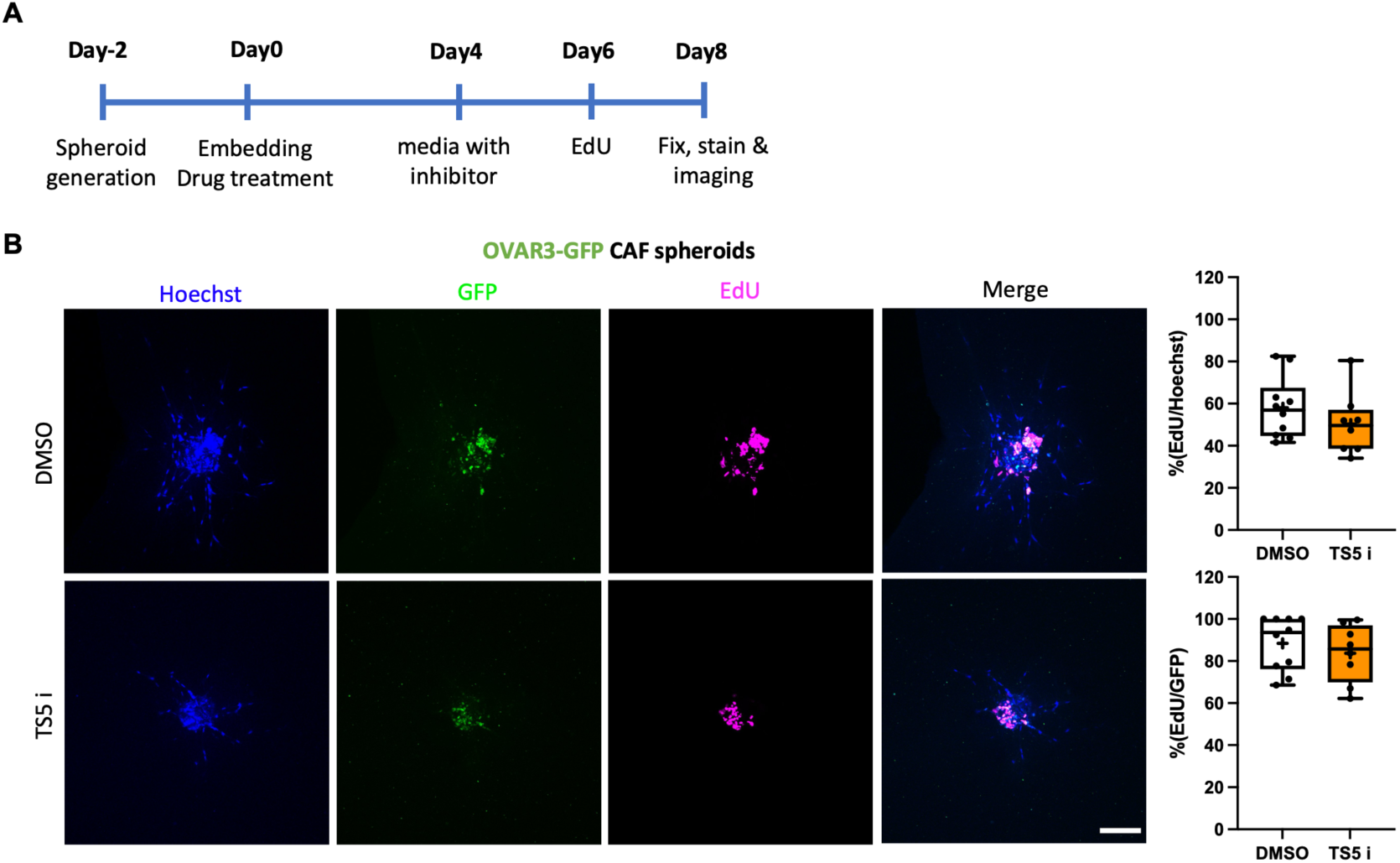
ADAMTSS inhibitor did not affect OVCAR3 cell proliferation in 3D. **(A)** Experimental timeline. **(B)** Spheroids generated with OVCAR3-GFP cells (green) and non-labelled CAFs (2:1 ratio) were embedded in 3mg/ml Geltrex, 3mg/ml collagen I and 25µg/ml fibronectin and cultured in the presence of DMSO or lOµM ADAMTSS inhibitor (TSS i). EdU was added at day 6, spheroids were fixed on day 8, stained with EdU detection reagent (magenta) and Hoechst 33342 (blue) and imaged with a Nikon Al confocal microscope. Scale bar, 200µM. The percentage of EdU positive cells was calculated against Hoechst or GFP. Data are presented as box and whisker plots (Min to Max,+ represents the mean) from N=3 independent experiments.

**Figure S7.**
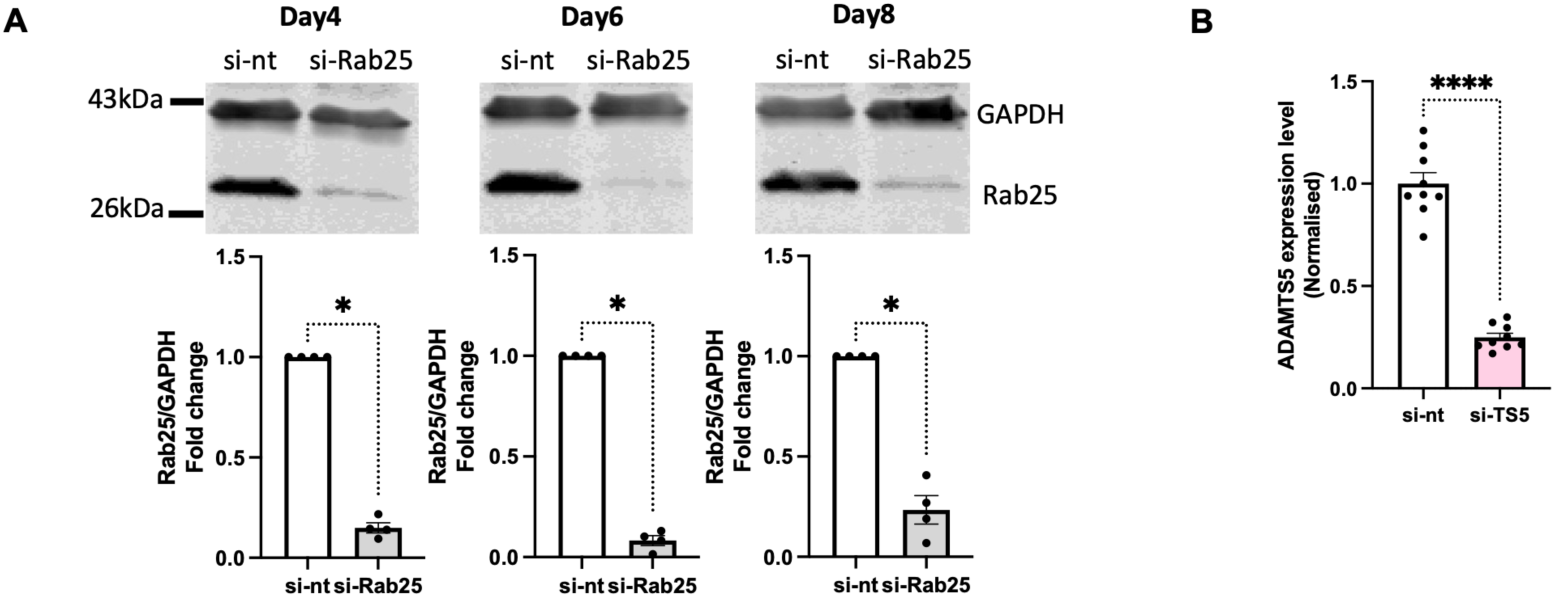
Rab25 and ADAMTSS KD efficiency in OVCAR3 cells. **(A)** OVCAR3 cells were transfected with a non-targeting (si-nt) or Rab25-targeting (si-Rab25) siRNA, cell lysates were collected after 4, 6 and 8 days and Rab25 and GAPDH protein levels were quantified by Western Blotting. Membranes were imaged with a Licor Odyssey Sa system, and the band intensity was quantified by Image Studio Lite software. Rab25/GAPDH intensity was normalised to si-nt. Data are presented as mean ± SEM from N=4 independent experiments; * p=0.0286, Mann-Whitney test. **(B)** OVCAR3 cells were transfected with a non-targeting siRNA control (si-nt) or ADAMTSS-targeting (si-TSS) siRNA and the mRNA levels of ADAMTSS and GAPDH were measured by qPCR. Data were normalised to si-nt and presented as mean ± SEM from N=3 independent experiments. ****p<0.0001, Mann-Whitney test.

